# Bimodally oriented cellulose fibers and reticulated homogalacturonan networks - A direct visualization of Allium cepa primary cell walls

**DOI:** 10.1101/2022.01.31.478342

**Authors:** William J Nicolas, Florian Fäßler, Przemysław Dutka, Florian KM Schur, Grant Jensen, Elliot Meyerowitz

## Abstract

One hallmark of plant cells is their pecto-cellulosic cell walls. They protect cells against the environment and high turgor and mediate morphogenesis through the dynamics of their mechanical and chemical properties. The walls are a complex polysaccharidic structure. Although their biochemical composition is well known, how the different components organize in the volume of the cell wall and interact with each other is not well understood and yet is key to the wall’s mechanical properties. To investigate the ultrastructure of the plant cell wall, we imaged the walls of onion (*Allium cepa*) bulbs in a near-native state via cryo-Focused Ion Beam milling (cryo-FIB-milling) and cryo-Electron Tomography (cryo-ET). This allowed the high-resolution visualization of cellulose fibers *in situ* (*in muro*). We reveal the coexistence of dense fiber fields bathed in a reticulated matrix we termed “meshing,” which is more abundant at the inner surface of the cell wall. The fibers adopted a regular bimodal angular distribution at all depths in the cell wall and bundled according to their orientation, creating layers within the cell wall. Concomitantly, employing homogalacturonan (HG)-specific enzymatic digestion, we observed changes in the meshing, suggesting that it is at least in part composed of HG pectins. We propose the following model for the construction of the abaxial epidermal primary cell wall: The cell deposits successive layers of cellulose fibers at −45° and +45° relative to the cell’s long axis and secretes the surrounding HG-rich meshing proximal to the plasma membrane, which then migrates to more distal regions of the cell wall.

## Introduction

Plants dominate the earth’s biomass^1^ and provide oxygen necessary for nearly all life on earth through photosynthesis. Photosynthesis allows the fixation of CO2 to form simple sugars through the Calvin-Benson cycle, breaking down water molecules and releasing oxygen^2^. A major fraction of the synthesized simple sugar is used to build up the plant pecto-cellulosic cell wall^3^. The cell wall is a heterogeneous mix of polysaccharides, mainly linear chains of *β*1-4-linked glucose (cellulose), pectins, which come in a wide chemical variety, and hemicelluloses, which are also chemically diverse^4,5^. The complex composite structure of the cell wall is crucial for shaping cells and, in turn, for cellular function. The unique feature of the cell wall in this context is its ability to resist chemical/enzymatic treatments and mechanical stress while still allowing cells to grow ^5^.

The major player in cell shape determination and cell wall stiffness is cellulose. Cellulosic glucan chains assemble to form higher-order fibers with amorphous and crystalline regions^6–8^.

The determinants of bundling of the cellulose fibers, mainly their interaction with hemicelluloses and pectins, are thought to be very important as they confer additional, higher-order mechanical properties^9,10^. The cellulose fibers are secreted into the cell wall by membrane-embedded hexameric Cellulose Synthase Complexes (CSC), each protomer, according to recent work, comprising a trimer of Cellulose Synthases (CESAs)^11,12^. In the current model, each CESA secretes a glucan chain, resulting in an elementary fibril secreted by a CSC that is composed of 18 glucan chains. It has been shown that the mature CSCs, upon delivery at the plasma membrane, associate with cortical microtubules *via* intermediary partners such as Cellulose Synthase Interactive protein-1 (CSI), which then guide the direction of cellulose synthesis *in muro*^13^. Although microtubule-guided cellulose synthesis is the most described and well-understood facet of this process, a microtubule-independent pathway has been characterized where CSCs separate from their microtubule track in favor of following an already existing cellulose fiber on the other side of the plasma membrane ^14^. The latter relies on integrating the newly synthesized fibers into an already existing bundle of microfibrils in the cell wall, also hinting towards a mechanism where the motile force of the CSCs is not cytoskeleton-dependent but rather propulsion due to cellulose crystallization^15^. A cohort of studies showed that the orientations of the cellulose fibers are consequential to the shape of a cell^16–19^ and the existence of a mechanical feedback loop where the cell is able to sense mechanical cues through its cortical microtubular network and adapt the cellulose fiber patterns in the cell wall ^20^.

While the cellulose fibers are thought to be the main load-bearing structures in the cell wall, pectins and hemicelluloses interact with them in ways still not fully understood. Hemicelluloses are currently hypothesized to tether cellulose bundles together and form load-bearing hotspots ^21,22^. Pectins, mainly homogalacturonans (HGs), making up to 60% of the dry weight of the primary cell wall ^4^, are hypothesized to surround all other components and act as a matrix ^8^. Composition, methylation state, and calcium levels have been shown to change the mechanical properties of pectins by altering the level of crosslinking ^23,24^.

Despite our knowledge of the chemical composition of the cell wall and of the diversity of the individual components, structural understanding of their secretion and interaction in the cell wall is underexplored. Cellulose-specific stains have been applied directly to live tissue to observe the cellulose fibers and follow their fate during cell elongation ^18,25^, but light microscopy does not offer the necessary resolving power to observe the cellulose fibers and their partners at nanometer resolution. White onion (*Allium cepa*) abaxial epidermal cell wall peels have been used in conjunction with high-resolution Atomic Force Microscopy (AFM) and field emission scanning electron microscopy to characterize the organization of the cell wall components at higher resolution ^9,20,22,26,27^. Despite the knowledge gained, AFM can only access the superficial layers of the cell wall leaving the rest of this polylamellate structure, estimated to be as much as 100 layers, unobserved. Having access to the depth of the cell wall allows a better structural understanding of the cell wall and its relation to cell shape. Here we used Cryo-Focused Ion Beam milling (cryo-FIB-milling) followed by cryo-Electron Tomography (cryo-ET) to observe plunge-frozen *Allium cepa* abaxial periclinal cell walls of onion scale epidermal cells throughout their depth, in near-native conditions.

The high-resolution data we gathered at multiple depths of the cell wall reveal the coexistence of cellulose fibers and a structure coined “meshing”, which our data suggest is made at least in part of homogalacturonan pectins. The fibers are shown to adopt a bimodular angular distribution creating layers of fibers of alternating angles of -/+ 45° relative to the cell’s long axis.

## Results

### Cryo-ET on epidermal cell wall peel lamellae allows the visualization of the plant cell wall in near-native conditions

White onion cell wall peels from the concentric scales, numbered #1 (outermost and oldest scale) inward to number 8 (innermost youngest scale), were generated as described previously (Figure 1A-C) ^28,29^. They were then flash-frozen and cryo-FIB milling was performed on the periclinal cell walls to produce lamellae ∼200nm in thickness, allowing access to the deeper layers (Figure 1D-H). As the angle of milling was well defined, it was possible to measure the depth of the tomograms in the cell wall (Figure 1I and J). Keeping in mind the known artifacts visible on the lamellae, such as curtaining, surface ice contamination, and surface gallium streaks (Figure 1K, red arrows, blue asterisks, and red arrowheads, respectively), tomographic data acquired in this way allowed visualization of the organization of the different elements in the cell wall at high resolution, in near-native conditions. Fields of fibers organized in arrays (Figure 1L, colored arrows) were observed, as well as small, intercalated patches of thin, reticulated densities we term “meshing” (Figure 1L, blue circles and Fig. 2). At the same time, we found the meshing to intercalate between bundles of cellulose fibers (Figure 2A-D, yellow and blue arrows and dashed line, respectively and supplemental video 1). Because manual segmentation of these two features was impossible, two Convolutional Neural Networks (CNN) were trained to recognize these two features using the EMAN2 software^30^. The fiber detection CNN, being very specific, yielded precise maps of the fibers (Figure S1A-C). However, the meshing detection neural network also detected the fiber densities in the tomogram. To circumvent this issue, subtraction of the CNN fiber map from the meshing-CNN map was performed, which assumes that all densities that are not fibers are associated to the novel meshing (Figure S2). The meshing is seen accumulating in patches between the fibers and connecting the fibers together. X-Z cross-sections of the segmentations corrected for lamella tilt allows qualitative assessment of the distribution of these two features within the volume (Figure 2E). In tomograms with a similar layout as in Figure 2A, the volume occupancy of the meshing versus that of the fibers ranged from 30% to 75% (Figure 2F).

**Figure 1.**
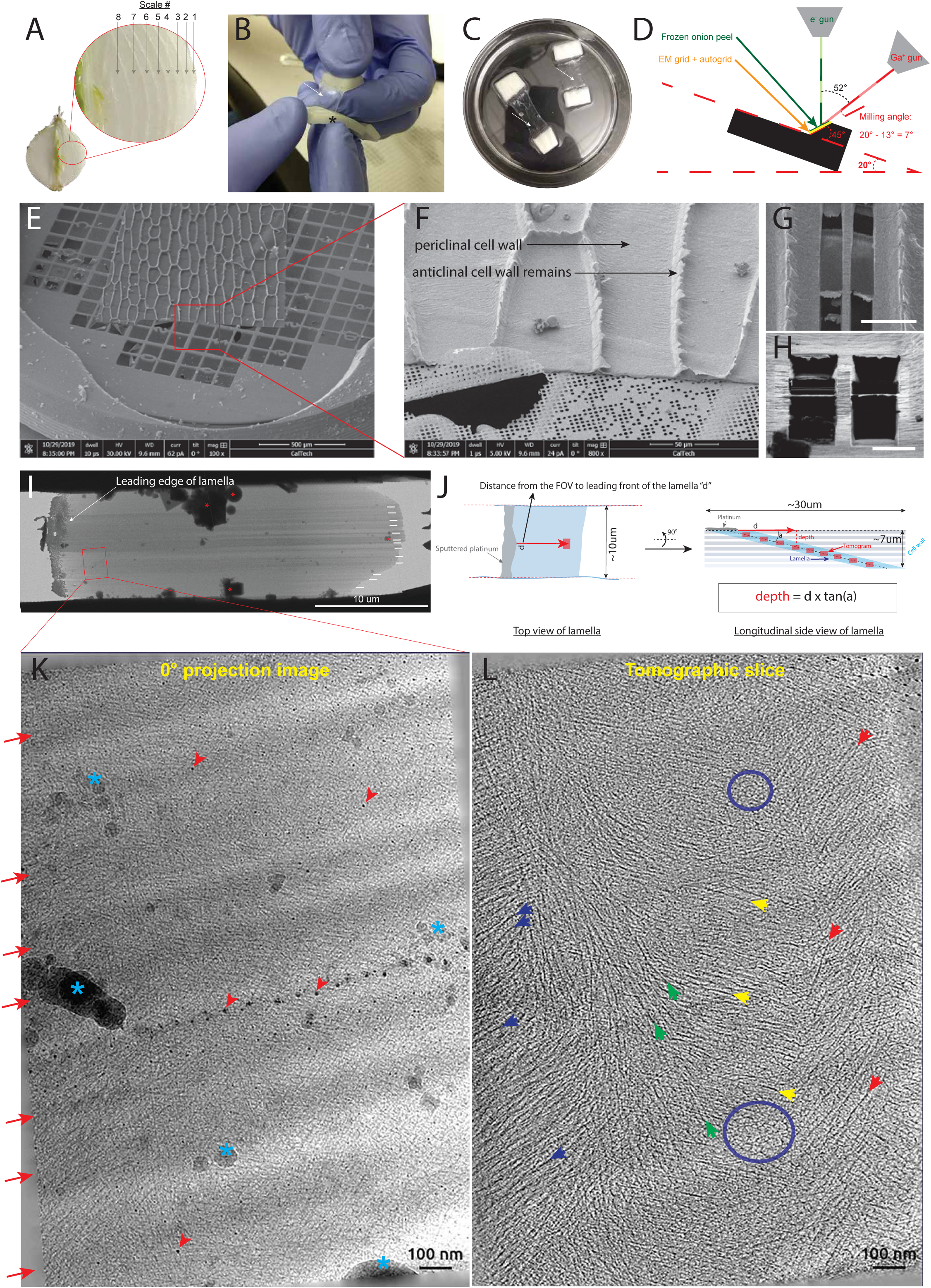
From fresh onion to reconstructed tomograms. (**A**) Half-cut onion showing the concentric scales. The inset shows the classic way the scales are numbered, from outermost to innermost. (**B**) Process of peeling the abaxial epidermal cell wall. (**C**) Cell wall peels (clear membranes) attached to the two thicker handles (white) incubating in HEPES. (**D**) Diagram of the SEM chamber and the position of the onion cell wall peel (green) relative to the FIB and electron beam. (**E**) SEM overview of a cell wall peel laid on an EM Quantifoil grid. (**F**) Magnified view of the red box in (E) showing the anticlinal and periclinal cell walls (where the milling was done). (**G**) SEM overview of two final lamellae milled in periclinal cell wall. (**H**) FIB view of the two same lamellae shown in (G). (**I**) TEM overview of a milled lamella. Curtaining is visible (white lines) and contamination is seen on the lamella (red asterisks). (**J**) Left, diagram of a lamella top view showing how the distance *d* from tomogram to leading edge of lamella is measured. Right, side view of a lamella illustrating how tomograms distributed along the length of the lamella can sample the different layers of the cell wall. (**K**) 0° projection image of the red boxed area in (I), 0.40 um below the surface of the cell wall in scale #6. Various typical FIB milling artefacts are visible: curtaining (red arrows), platinum projections (red arrowheads), ice contamination (blue asterisks). (**L**) Central tomographic slice of the same area shown in (K). Numerous fibers are visible (blue, green, yellow, and red arrows) and small patches of short rod-like, branched densities can be seen between the fibers (blue circle).

**Figure 2.**
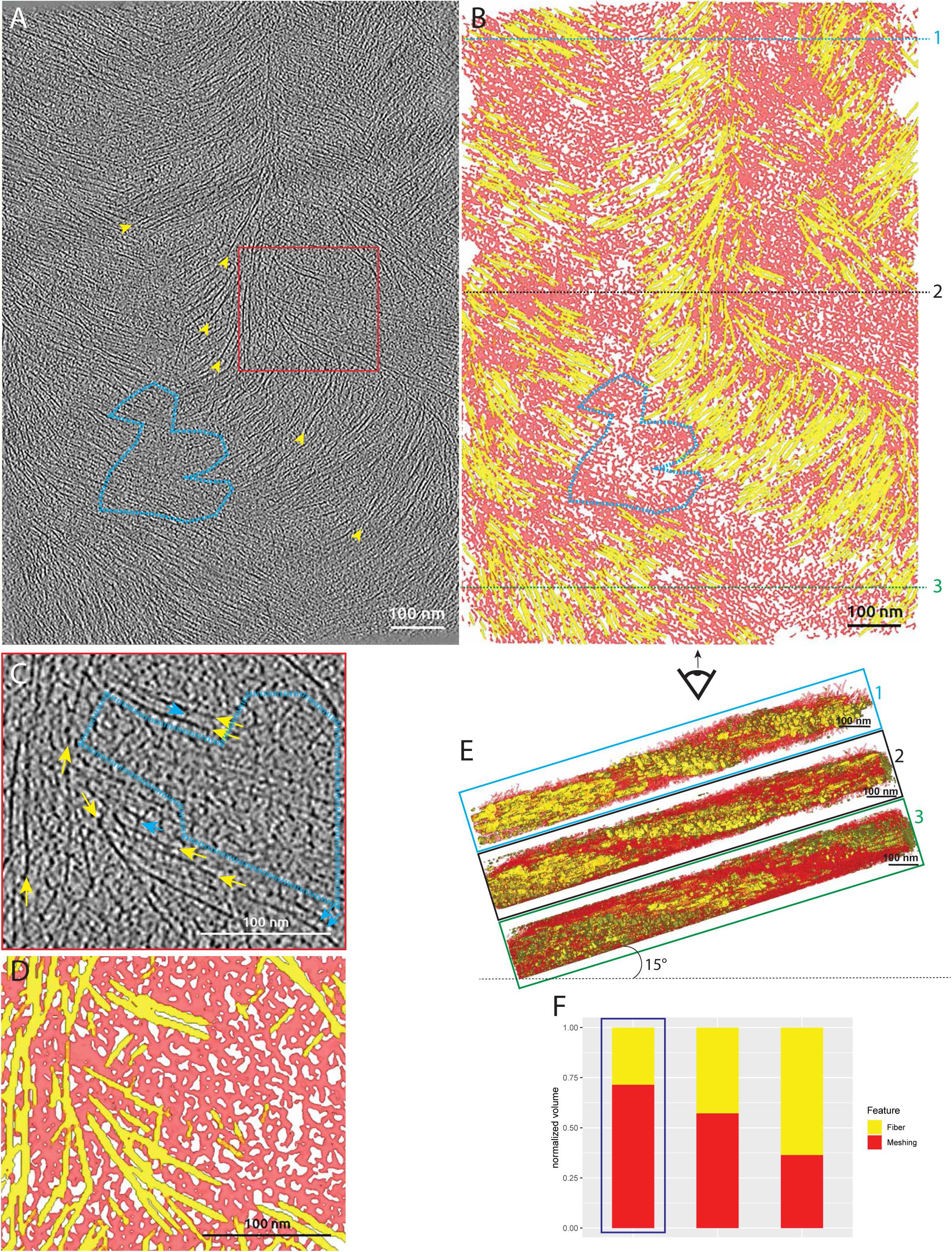
Two features in tomograms of onion cell walls. (**A**) Tomographic slice showing fibers (yellow arrowheads) and the filling material in between the fibers (blue circles), the meshing. This tomogram originates from scale #6, 0.64 um below the surface of the cell wall. (**B**) CNN segmentations of the fibers (yellow) and the meshing (red). (**C**) Magnified view from the red boxed region (A) showing fibers (yellow arrows), tethers between the fibers (blue arrows), and patches of meshing (blue dashed region). (**D**) CNN segmentation of the magnified region. (**E**) Transverse views at different Y-levels of the segmented volume shown in (B). Tilting is correction for the inclination of the lamella relative to that of the wall. Alternations of fibers (yellow) and meshing (red) are observed. (**F**) Relative occupancy of fibers vs. meshing in 3 tomographic volumes equivalent to the one shown in this figure (blue squared column is the tomogram shown).

### Fibers travel straight and horizontally in the cell wall and adopt a bimodal angular distribution

To produce a vector representation of the fibers suitable for geometrical analysis, a template-matching strategy using the Amira TraceX add-on^31^ was used on the fiber-CNN maps (Figure S1C-F). The following results were extracted from a total dataset of 31 tomograms acquired across the onion scales #2, 5, 6, and 8 (Figure 1A, see supplemental table 1 for a precise description of the data and samples). The density distribution of the orientation of the fibers was analyzed for each tomogram. 26 out of the 31 tomograms considered showed a bimodal distribution (Figure 3A and B, supplemental video 2), the 5 others exhibited a unimodal distribution (Figure S3D). All the tomograms displaying a bimodal distribution had very similar angles to the long cell axis, averaging 42° ± 8° (n = 31 tomograms) and 135° ± 10° (n = 26 tomograms), showing a difference between the two modes of ∼90°. Since all angles were calculated clockwise, the 135° relative to the cell’s long axis is equivalent to a 45° angle counterclockwise (Figure 3B and C). When organized according to the scale number where the tomogram was acquired, the density distributions show very similar modes (Figure 3C), suggesting that this bimodal distribution of the orientation of the fibers is consistent throughout all developmental stages studied. Fibers with the same angle cluster together according to their Z-height within the tomographic volume (Figure 3D), creating horizontal layers of cellulose fibers alternating between 45° clockwise/counterclockwise (Figure 3E).

**Figure 3.**
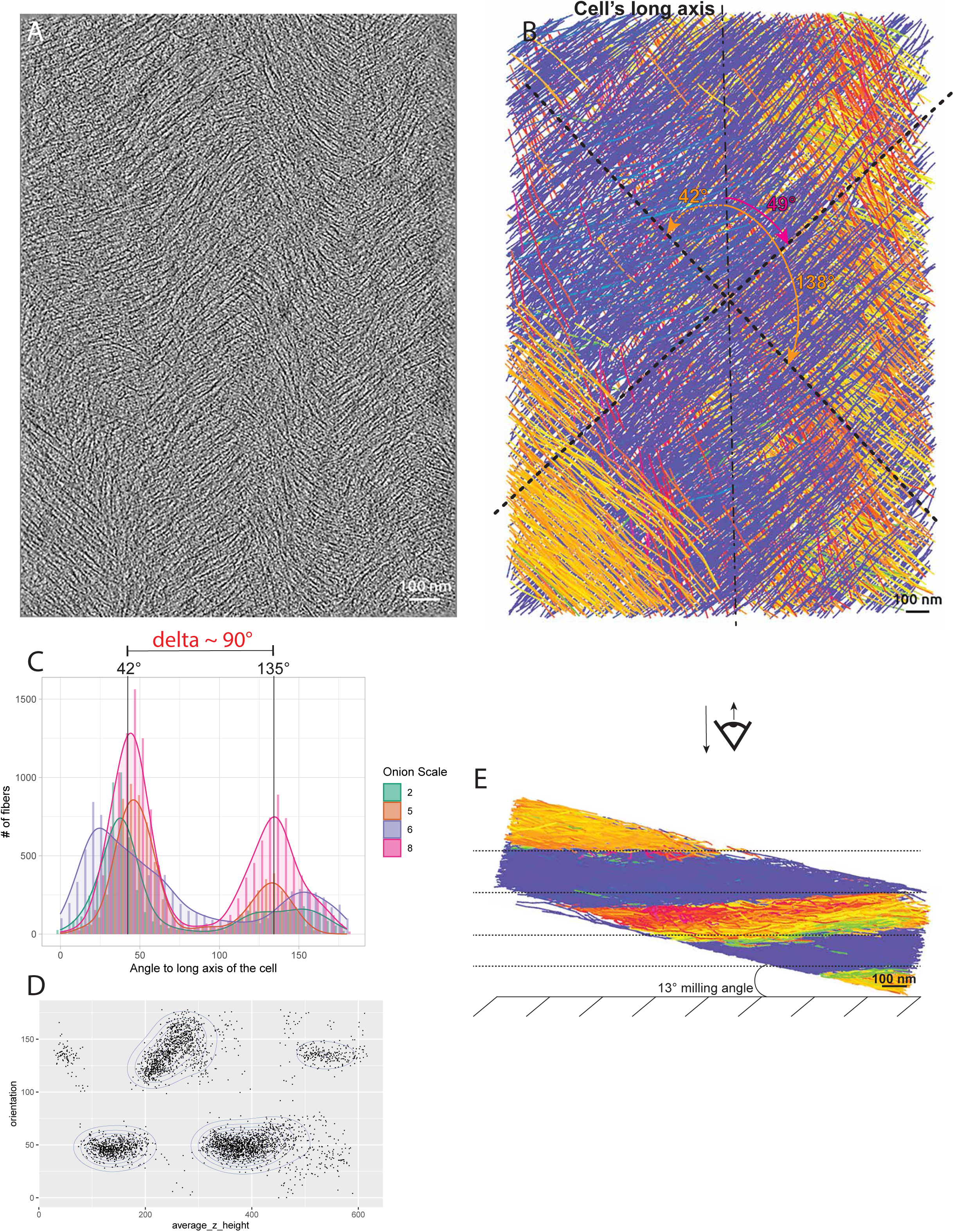
Cellulose fibers organize in a bimodal angular pattern. (**A**) Tomographic slice of the cell wall from scale #8, 3.36 um below the surface. (**B**) Automated segmentation of the volume shown in (A). Color coding is according to the clockwise angle of the fiber relative to the cell’s long axis (∼ vertical dashed line). The two dashed crossed lines indicate the two main angular modes in this volume. They are 49° and 138°. The latter is equivalent to a 42° counterclockwise angle. (**C**) Distribution plot of the angle of the fibers relative to the cell’s long axis by scale number. The 42° and 135° angles correspond to the global modes, aggregating all fibers from all scales. The difference between these two modes is ∼90°. (**D**) Scatterplot of angles of fibers vs. their average height in the tomographic volume shown in (A) and (B). (**E**) Bottom cross-sectional view of the segmented volume shown in (B).

Three more distribution patterns were observed in addition to a perfectly staggered pattern (Figure 3D and Figure S3A): Overlapped (12 out of 31 tomograms), with fibers of both modal angles mixed at all heights of the tomographic volume (Figure S3B). Staggered-overlapped (9 out of 31 tomograms), similar to the staggered pattern but with overlapping (Figure S3C). Unimodal (5 out of 31 tomograms), with only one modal angle, (Figure S3D).

The effect of the depth in the cell wall and the aspect ratio of the cell on the angular distribution was investigated on a per-scale basis. The bimodal angular pattern is found throughout all scales studied (#2, 5, 6, and 8), at all depths and cell aspect ratios where data were acquired (Figure S4A-C). The aspect ratios of the milled cells fell into the ranges measured from the light microscopy montages. The average aspect ratios for scales 2, 5, and 8 and their standard deviation overlap strongly (4.3 ± 1.9, 3.8 ± 1.5, and 4.1 ± 1.7, respectively), suggesting that there is little to no change in the cell’s aspect ratio as the scale is pushed outward during growth of the onion (Figure S4D and E).

The straightness and the horizontality (relative to the horizontal plane of the cell wall) of each fiber were also analyzed by computing the average radius of curvature and average slope of each fiber, respectively (see methods for details on the computation of these parameters). The average radius of curvature measured throughout all the tomograms is 225 ± 90 nm, which suggests that the fibers are overall straight (Figure S5A and B). The average slope measured throughout all the tomograms is 0.02 ± 0.4 and is centered around 0 (Figure S5C), indicating that the fibers describe horizontal trajectories within the volume of the wall, clearly observable when looking at cross-sections in the segmentations (Figure S5D and E).

In summary, these results show that the fibers organize in layers that alternate between −45°/45° relative to the cell’s long axis, are relatively straight and travel horizontally relative to the cell wall horizontal plane.

### The meshing accumulates at the surface of the cell wall

We also characterized the meshing, which takes the form of thin and short fibrous densities that either reticulate forming a web-like network (Figure 4A and B) or tether cellulose fibers together (Figure 4C). Tomograms acquired proximal to the platinum layer and thus close to the previous wall interface with the plasma membrane (Figure 4D and J) show extended areas of reticulated meshing (Figure 4E, F, H, and I and Figure 4K, L, N, and O, red and black dashed delineations) accompanied by a reduction in the concentration of fibers. In regions of enriched meshing, the relative volume of the wall region manifesting mesh can be above 50% (Figure 4G and M). Having lamellae milled at an angle allowed probing the structure of the cell wall not only proximal to the cell surface but also more distally in the cell wall (Figure 5A). We were therefore able to follow the distribution of the meshing within the depth of the cell wall. Proximal to the cell wall inner surface, transition areas could be observed even within a single tomographic volume. A sub-region of the tomogram was depleted in meshing (Figure 5B, left of the yellow dashed line and supplemental video 3) and contained ordered bundles of fibers, while the other sub-region was enriched in meshing (Figure 5B, D and E and supplemental video 3) and exhibited more disordered arrays of fibers. In contrast, tomograms acquired deeper in the cell wall (farther from the plasma membrane) had reduced amounts of meshing and displayed fibers with an increased degree of bundling and order (Figure 5C, F and G and supplemental video 3). Quantitative analysis of the segmented meshing volume to segmented fiber volume ratio shows a gradual decrease in the amount of meshing as a function of depth in the cell wall (Figure 5H).

**Figure 4.**
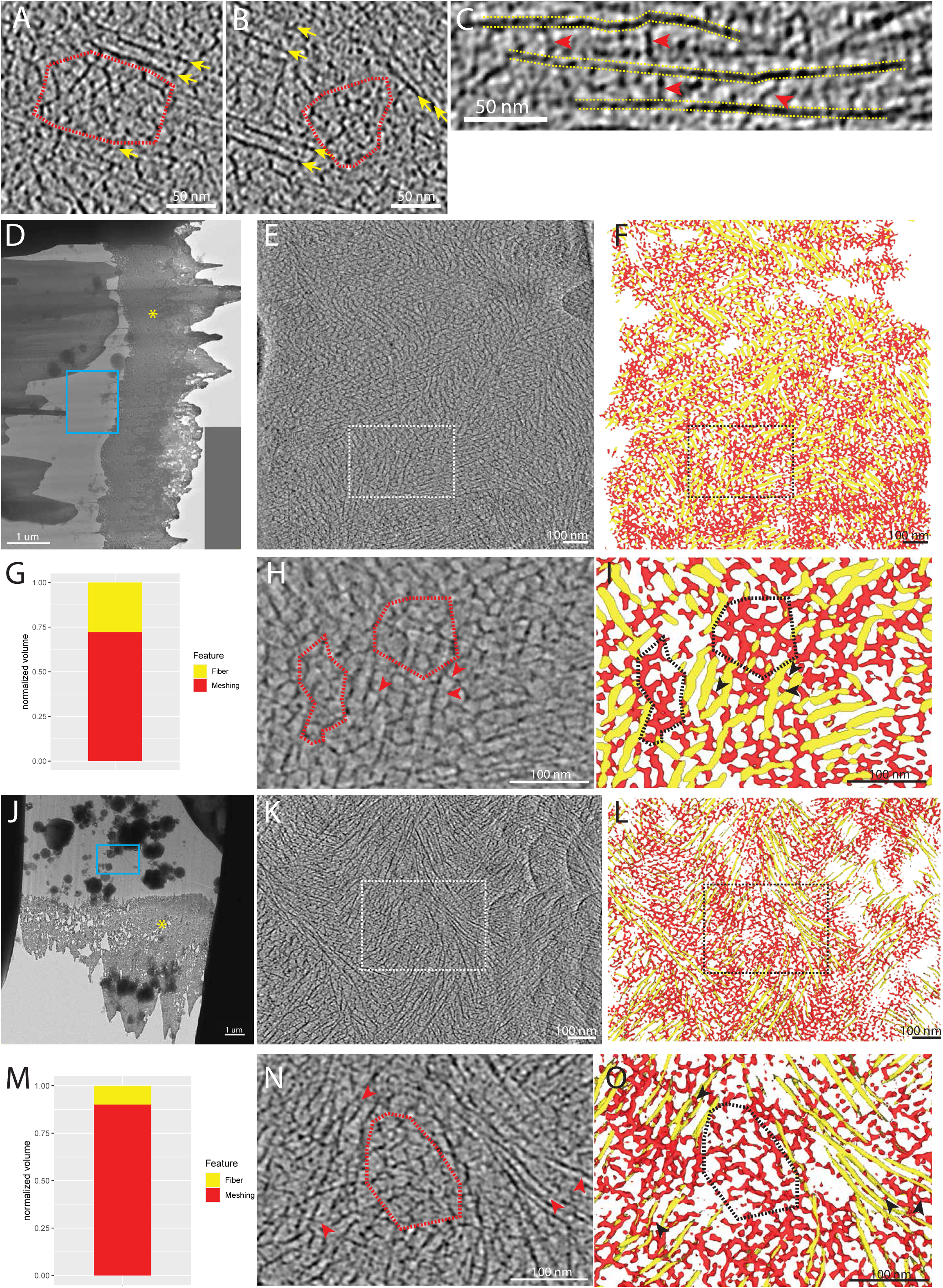
The meshing is seen in patches and also tethering the fibers together. (**A, B**) Examples of small patches of meshing (circled in red) surrounded by fibers (yellow arrows). The meshing is characterized by small, branched segments with no particular orientation, creating a reticulated network. (**C**) Examples of meshing segments (red arrows) tethering fibers together (yellow dashed lines). (**D, J**) Overviews of lamellae. Yellow asterisk points to the platinum layer. (**E, K**) Tomographic slices of tomograms acquired near the top of the cell wall (blue rectangle in (D) and (J), respectively) at 0.15 um and 0.55 um below the surface, respectively. These tomograms are enriched in meshing as many reticulations can be seen. (**F, L**) Associated segmentation of the tomographic slice shown in (E) and (K), respectively. Meshing is in red and fibers in yellow. (**G, M**) Relative quantity of meshing vs. fibers in the tomogram shown in (E) and (K), respectively. (**H, N**) Magnified views of the white rectangles shown in (E) and (K), respectively. Examples of patches of meshing are shown (red dashed circles) and events of fiber tethering are highlighted (red arrowheads). (**I, O**) Corresponding segmentation of the magnified view (H) and (N), respectively.

**Figure 5.**
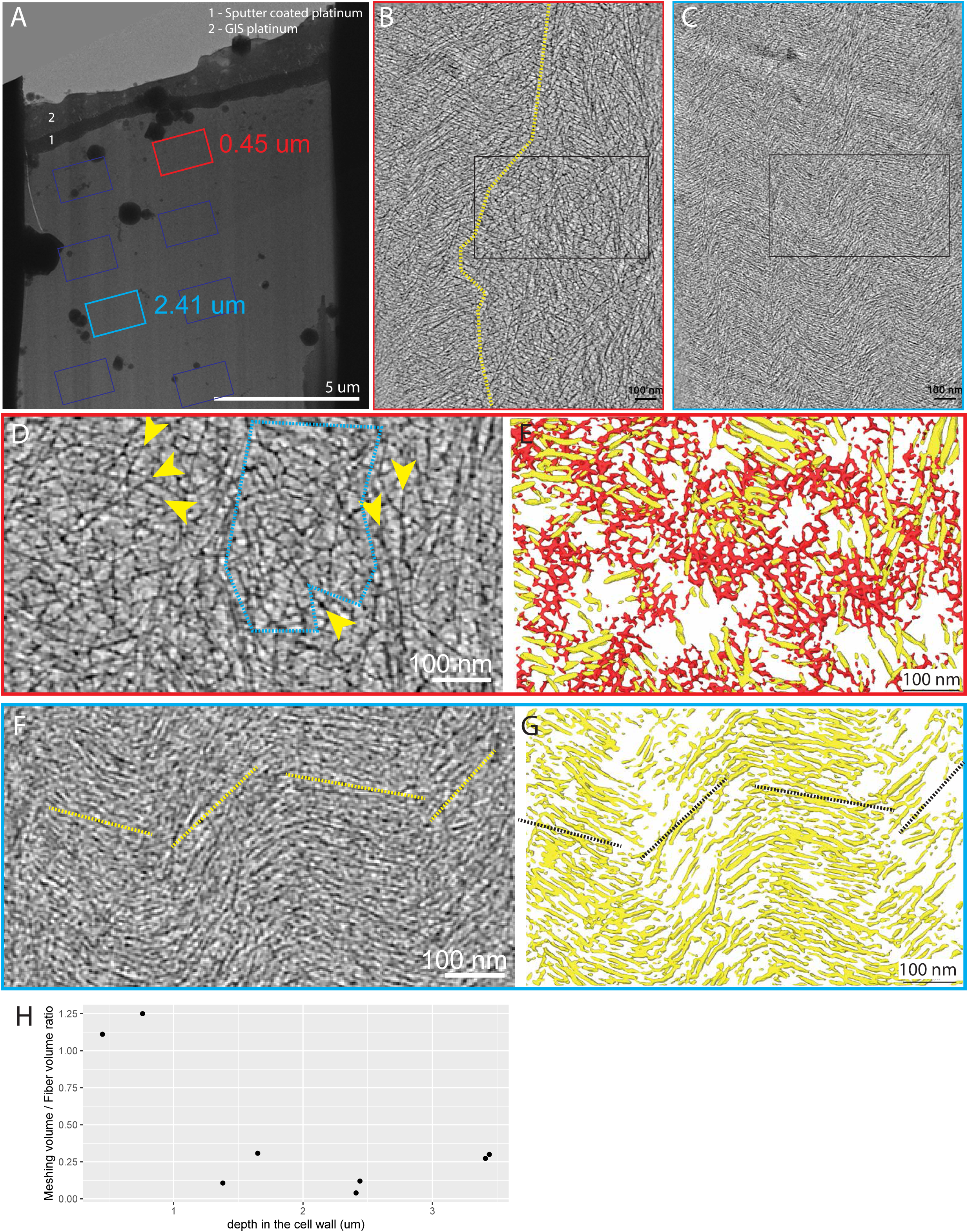
The meshing is concentrated at the top of the cell wall. (**A**) Overview of lamella milled in a non-treated cell wall peel from scale #2. Rectangles show where tilt series were acquired. (**B**) Tomographic slice acquired near the top of the cell wall (red rectangle in (A), at 0.45μm from the surface). The dashed yellow line indicates the visual limit between an area with a loose network of fibers with a substantial amount of meshing intercalated between the fibers (right of the line), and an area where the fibers seem more bundled together and less meshing is visible (left of the yellow line). (**C**) Tomographic slice acquired further down the cell wall (blue rectangle in (A), at 2.41μm from the surface). It shows a denser network of fibers with virtually no visible meshing. (**D**) Magnified view from the black rectangle in (B). Extensive patches of meshing intercalated with fibers can be observed (blue dashed circle and yellow arrows, respectively). (**E**) Associated segmentation of the magnified view (D). (**F**) Magnified view from the black rectangle in (C). Tightly packed fibers with very constant orientations are seen. Yellow dashed lines show the general orientation of the layers visible in this tomographic slice. (**G**) Associated segmentation of the magnified view (F). (**H**) Meshing vs. fiber volume ratio calculated from the CNN segmentations in the 8 tomograms extracted from this lamella. The ratios are plotted against the depth from at the tomograms were extracted from the cell wall.

Taken together, these results suggest the meshing is secreted out of the cell, accumulates at the cell wall-PM interface, and reticulates between the fibers of the first layers of the cell wall.

### Enzymatic digestion of the homogalacturonans alters the morphology and abundance of the meshing

We sought to identify the chemical nature of the meshing. According to the literature, pectic homogalacturonans (HGs) make up 50% of the primary onion cell wall ^32^. In the most recent models of the interactions between the different components of the primary cell wall, pectins are thought to surround and tether the cellulose fibers in a calcium-dependent manner ^8,10,23^. Considering this, cell wall peels treated with either BAPTA, a calcium chelator, or *Aspergillus* Pectate Lyase (PL), an enzyme that digests de-methylesterified homogalacturonans, were processed by cryo-FIB milling and cryo-ET to see whether the previously observed meshing would be morphologically altered. The efficiency of the treatments was verified by staining treated and non-treated peels with Chitosan OligoSaccharide Alexa-488 (COS488), an HG-specific fluorescent probe (Figure S6A and B). The decrease in fluorescence (most apparent in the PL-treated material) suggests that these treatments reduce the pectin content in the peels (Figure S6C-F). While applying the onion peels to the EM grids, we noticed that the PL-treated ones seemed to exhibit greatly reduced stifness. Cryo-SEM images showed a clear difference in the thickness of the peels and a qualitative reduction in the prominence of the bases of the torn-off anticlinal cell walls, suggesting that the specific digestion of demethylated pectins from the cell wall affects cell wall thickness and the continuity between periclinal and anticlinal cell walls (Figure S7).

BAPTA-treated cell wall peels show visible meshing, as in untreated walls in increased concentration proximal to the leading edge of the lamella (Figure 6A, C and D, red arrowheads). PL-treated cell wall peels show no meshing at all or remnant densities between the fibers and in small patches that we interpret as partly digested meshing (Figure 6B, E and F). Quantification of meshing volume versus fiber volume as a function of depth of the tomogram in the wall in the non-treated condition clearly shows a gradual decrease (Figure 6G, green dots). The unusually elevated ratio (∼9 fold more meshing, Figure 6G black arrow) represented a region of the cell wall ∼500nm below the surface (tomogram shown in Figure 4J-O) and was excluded from the computation of the average. The ratios found at the surface of the cell wall in the BAPTA-treated peels show a steady amount of meshing, overall lower than in the same non-treated regions of the cell wall (Figure 6G inset, average ratios of 0.82 ± 0.54 and 1.4 ± 2.2 for BAPTA and non-treated, respectively). In the PL-treated peels, the ratios were much lower (Figure 6G, inset, 0.26 ± 0.27). This suggests that PL treatment reduces the amount of meshing and alters its morphology, indeed in some cases making it practically disappear. Angular distribution of fibers was also assessed in the BAPTA-/PL-treated cell wall peels and the bimodal angular distribution pattern was conserved (Figure 6H).

**Figure 6.**
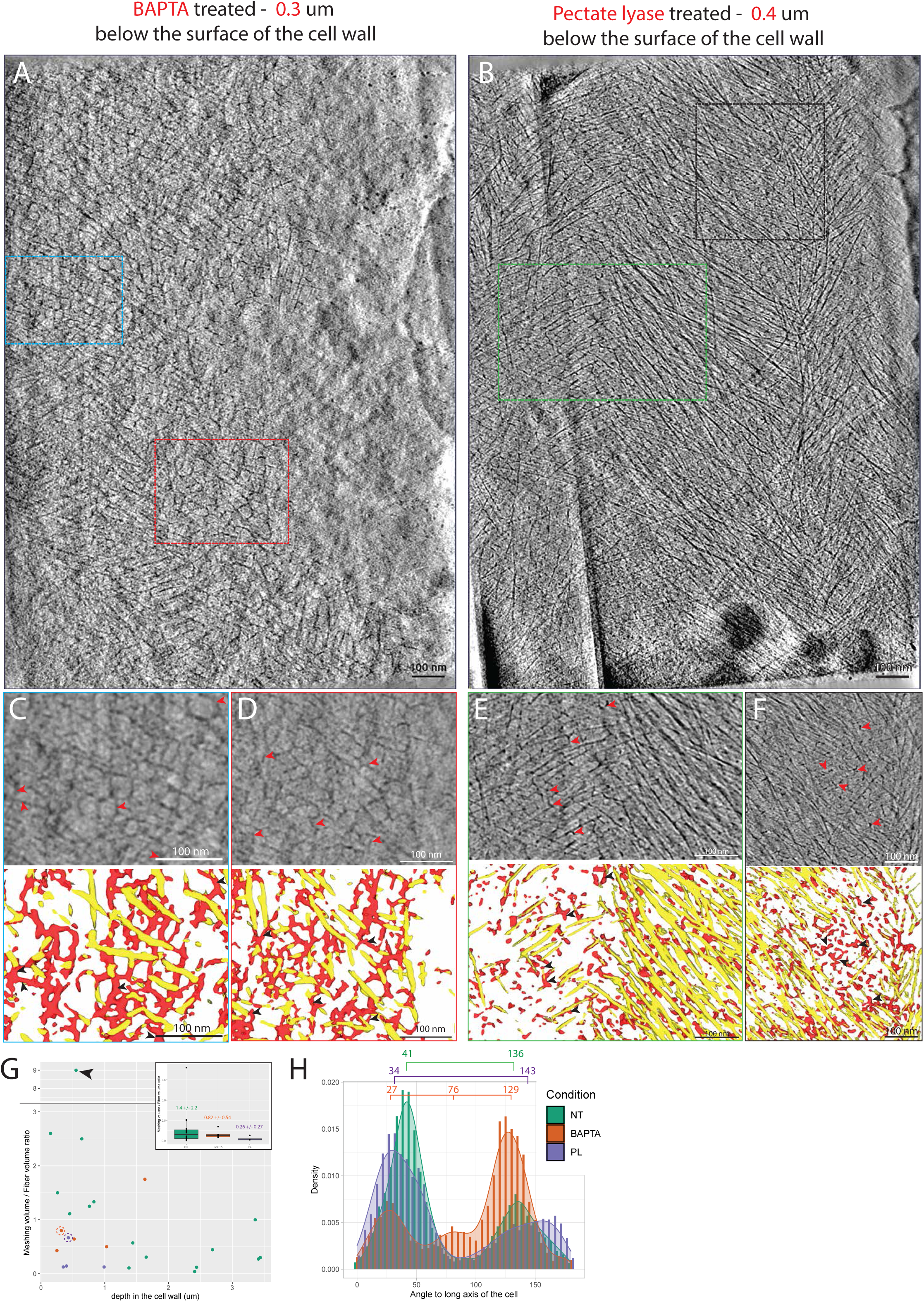
The morphology of the meshing is affected by pectate lyase but not by BAPTA. (**A**) Tomographic slice 0.3 um under the surface of a BAPTA treated cell wall. Meshing patches can be seen among the fibers. (**B**) Tomographic slice 0.4 um below the surface of a PL treated cell wall. Meshing remnants can be seen around the fibers. (**C, D**) Magnified views of areas of the tomogram (blue and red rectangles in tomogram (A), respectively) displaying meshing patches (red arrowheads) with the associated segmentations. (**E, F**) Magnified views of areas of the tomogram (green and black rectangles in tomogram (B), respectively) displaying small remnant densities in between the fibers (red arrowheads) with the associated segmentations. (**G**) Meshing vs. fiber volume ratio calculated from the CNN segmentations calculated from 16, 5, and 4 tomograms from non-treated, BAPTA and PL treated peels, respectively. Black arrow points to the unusually high meshing/fiber ratio. The inset boxplot shows the mean meshing/fiber ratio for each condition. The orange and purple dashed circles indicate the data points linked to tomograms shown in (A) and (B), respectively. (**H**) Distribution plot of the angle of the fibers relative to the cell’s long axis by condition. The brackets show the modal values for each of these conditions.

To assess whether the treatments had an impact on the cellulose fiber diameter, averages were generated for each condition (Figure 7A-C) and their cross-sectional thickness were compared by calculating the Full-Width-at-Half-Maximum (FWHM) on the full-length average density profiles. We were not able to measure a significant difference between the three averages generated (5.3, 6.0 and 6.3 nm cross-sectional thicknesses for the non-treated, BAPTA and PL conditions, respectively) (Figure 7D), suggesting the treatments did not alter the diameter of the cellulose fibers.

**Figure 7.**
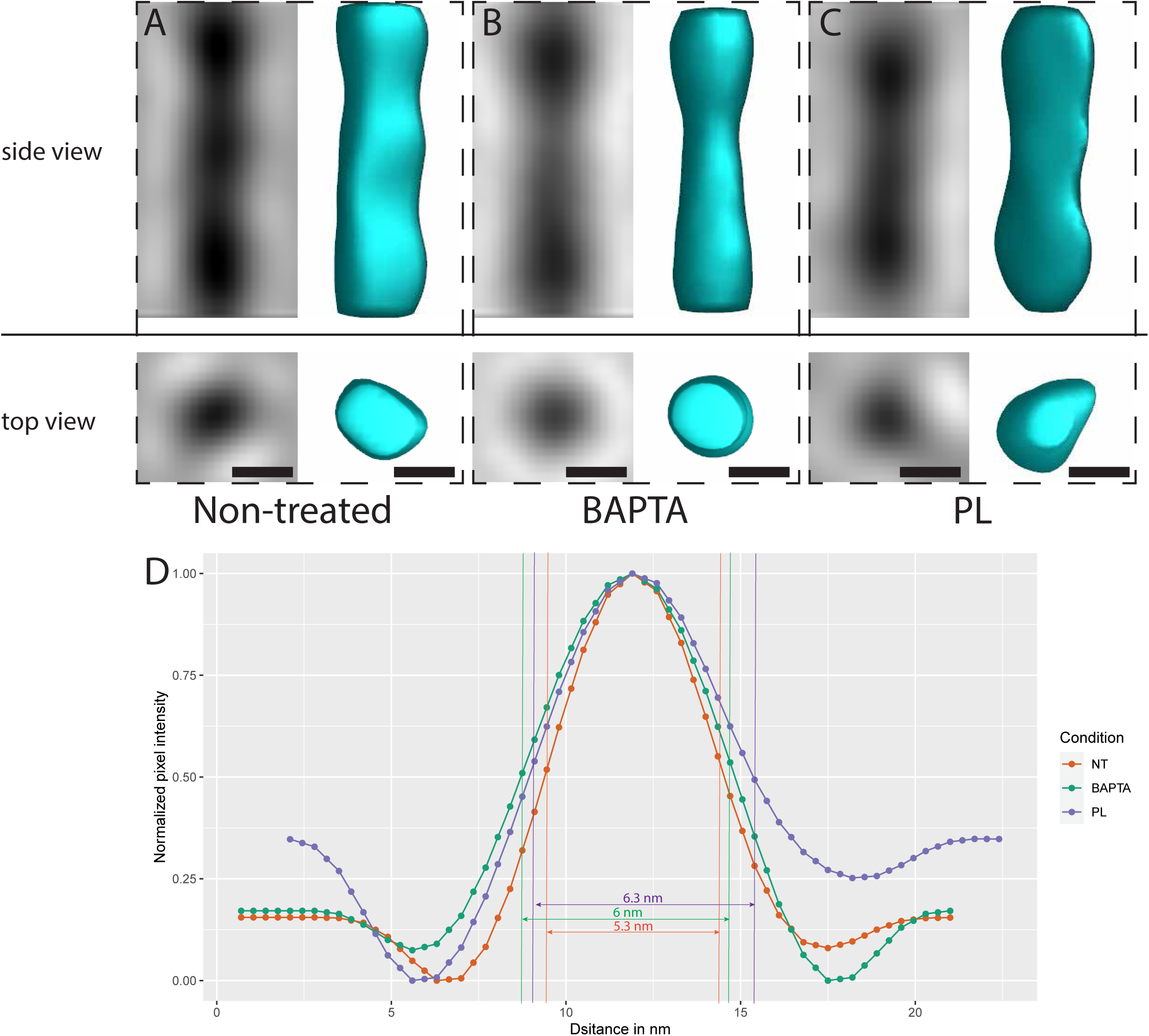
Fiber averages in non-treated and treated conditions. **(A-C**) Side views (top panels) and top views (bottom panels) of the fiber averages for the non-treated, BAPTA and PL condition, respectively. Scale bar = 5 nm. (**D**) Sideview profiles of the averages shown in (A-C) and the FWHM measurements.

### Purified pectins reproduce the morphology of wall meshing

To test our hypothesis that this meshing network seen around the fibers in the tomograms, altered in the presence of PL, is made of HGs, we imaged purified pectins in aqueous solution. Citrus pectins with an 89% content in galacturonic acid (HG) and a degree of methylation of 38% were observed using cryo-ET. As a negative control, solvent only (DI water) grids were also prepared.

The latter showed no features (Figure S8A), while the HG solution showed reticulated networks reminiscent of the meshing seen in the native cell walls (Figure S8B).

## Discussion

### Implications of the bimodal angular distribution

A hallmark observation in this work is the remarkable alternation of the cellulose layers between ±45° relative to the cell’s long axis (90° total angle between successive cellulose layers, supplemental video 2), confirming the crossed-polylamellate organization of the onion primary cell wall ^22,26^. Previous AFM data has observed a similar pattern at the inner surface of the cell walls but only in younger scales (8^th^)^28^. Along with the measurement of the aspect ratios at the different scales, Kafle et al. suggest that the anticlinal abaxial epidermal cells, as they expand and the scale ages, gradually shift from laying down cellulose fibers with no modal orientation to laying down the fibers in two modal orientations (±45° from the longitudinal axis of the cell), and finally in one modal orientation (90°), therefore linking the aspect ratio of the cell directly to the orientation of the fibers in the inner layers of the cell wall. Our observations show that the ±45 bimodal angular distribution is found ubiquitously at all the scales studied (Figure 3C), at all depths of the cell wall and all aspect ratios of the cells observed (Figure S4A and B).

Additionally, our light microscope observations of cell wall peels from different scales (Figure S4D and E) show different average aspect ratios and distributions than reported in Kafle et al. 2013 (aspect ratios for scales 2, 5, 8, and 11 of 5.2 ± 0.4, 2.6 ± 0.1, 3.4 ± 0.1 and 2.9 ± 0.1, respectively). This difference can be explained by how the analysis was conducted. Where we performed the measurements on several thousands of cells (Figure S9) in 3 independent onions, Kafle et al. measured the aspect ratio in ∼100 manually selected cells, potentially leading to more bias. There may also be a difference in the onions used, as the source was not genetically characterized in both cases.

Our ability to reach the deeper layers of the cell wall allowed observation of the bimodal angular distribution pattern throughout the whole depth of the cell wall (Figure S4A), suggesting there is no gradual reorientation of the cellulose layers as the cell expands, contrary to what has been observed in *Arabidopsis thaliana* root epidermal cells from the elongation zone ^18^ - as might be expected of cells where the aspect ratio does not substantially change with growth.

### Layering patterns

The case of layers of cellulose fibers stacked on each other, which we termed “staggered” (5/31 tomograms, Figure 3D and Figure S3A), was not the only one encountered. “Overlapped” instances (12/31 tomograms, Figure S3B) where the layers were fully intertwined with each other, and “Overlapped/staggered” instances (9/31 tomograms, Figure S3C) where the layers were stacked but were also partially overlapping suggests that multiple factors influence the trajectories of the Cellulose Synthase Complexes (CSCs).

In the light of our observations, we suggest that microtubule guidance can initiate a new layer with a different orientation from the previous one, which can then be maintained and reinforced by the microtubule-independent cellulose guidance acting as a positive feedback mechanism ^14^. Creating cleanly staggered alternating layers of cellulose ±90° from each other would require very drastic switches between these two guiding modes.

This can either be done by a complete synchronous turn-over of the present CSCs in the plasma membrane, which has been measured to require 8 minutes ^33^, a cessation of cellulose synthase activity that is known to be regulated through phosphorylation ^34,35^ or a sudden upregulation of CCs and CSI1 translation or delivery to the plasma membrane to redirect the active CSCs to reattach to microtubules so as to create a layer at a new orientation. It remains to be determined how the angles of −45° and +45° are conserved throughout the cell wall. Overlapped events where a mix of orientations are observed at a given height may indicate areas of the cell wall where this sudden directional switch did not occur or failed, hence the gradual transition. These events are the majority (21 if combining “overlapped” and “overlapped/staggered”), indicating that this is the typical mode of orientation switching. “Monolayered” instances (5/31 tomograms, Figure S3D) highlight areas where the cellulose layers are too thick to be entirely captured in one tomographic volume. Given their frequency, we postulate that this is not the norm. According to our model, an uninterrupted positive feedback or a long-term switch delay could create such layers. What regulates this switch and when it occurs is unknown.

### The mechanical relevance of a long-range straight fiber structure

Our results indicate that the cellulose fibers are generally straight with a mean radius of curvature of 225 ± 90 nm though allowing some bend with the curvature radii of individual fibers being as low as ∼50nm (Figure S5A and B). Live imaging of GFP-tagged CSCs in the membrane in Arabidopsis leaf cells showed turning angles ranging from 10° to 90° trajectories ^14^. Additionally, AFM assays coupled with stretching of onion cell wall peels have shown how some fibers are able to form local high curvatures and kinks under particular mechanical constraints ^22^ while the surrounding ones stay straight. We did not spot kinking events in the raw tomograms but cannot dismiss them as the volumes contained high numbers of fibers. Our segmentation method, using conjunctly CNNs and template matching, turns out to be very efficient for relatively straight fibers. However, we do not believe our method of quantifying the radii of curvature allowed recognition of local kinking events for two reasons:

i. we measured the average radius of curvatures of the whole fibers, masking local kinks due to averaging.
ii. The search cone used by Amira to trace the fibers on the CNN-segmented fiber maps had an angle of 37°, thus limiting its search to tracing within that range, excluding any fibers with kinks >37°.
iii. The majority of the fibers with low radii of curvature in our segmented volumes were actually tracing artefacts where two neighboring or crossing fibers with different orientations were erroneously connected.

It is known that the para-crystalline cellulose fibers can have fluctuating degrees of crystallinity along their length, with regions more crystalline or more amorphous than others. Regions with high crystallinity are associated with mechanical stiffness and, therefore, straightness, while amorphous regions are thought to be more flexible ^36^. The local low bending radii observed in our data and the local kinks observed in previous work ^22^ could correspond to regions of relatively lower crystallinity with an increased amount of amorphous cellulose. Regulating the crystallinity of the cellulose could be an additional means to regulate cellulose fiber stifness.

The latest molecular dynamics model of the cell wall, encompassing prior AFM work, predicts much more cross-linking between the cellulose fibers than observed in our tomograms ^37^, although the AFM work does not distinguish if crossing fibers are interacting or just intersecting on slightly different Z-levels ^22,26,28^. Our cryo-ET reconstructions seem to indicate that the fibers within a layer do not interconnect with each other but rather bundle together in a parallel fashion, tethered by the surrounding meshing (Figure 4).

Because of the ability to reposition our volumes in 3-dimensions, we were able to assess how the fibers run horizontally relative to the plane of the cell wall by calculating the slope of the fibers within the volume, 0.02 ± 0.4 on average (Figure S5C-E). Although this is the general assumption, it is to be emphasized that the cellulose fiber most likely comes out of the CSC orthogonally to the plasma-membrane, which means that a redirection of the nascent fiber is necessary for its integration in the cell wall. The movement of the CSCs caused by the crystallization of the fiber ^15^ coupled with the pressure exerted from the existing cell wall which was shown to guide crystallization ^14^ can explain the flattening of the fiber orientation and its integration in the cell wall. A similar process is at play in the building of the crystalline-cellulose ribbons in the *Gluconacetobacter hansenii* bacterium, where the proper crystallization of the nascent cellulose fibers requires the proximity of the existing extracellular ribbon ^38^.

### The meshing gradient and the mechanical properties of the cell wall

Tomograms of the near-native cell wall show an interconnected network of meshing and cellulose fibers, confirming the single network model of the primary cell wall ^39,40^. In some instances, overlaying the fiber and meshing segmentations shows clear layering (Figure 2E, supplemental video 1) of the two features. This meshing is therefore not just located at the surface of the cell wall but seems to serve as a matrix between the fibers even in more distal layers of the cell wall. This layering can be achieved by two means: i) alternation of secretion of a meshing and cellulose layer, or ii) the meshing is secreted at a constant rate and the bundling of the cellulose fibers as they are laid down gradually excludes or squeezes the greater part of the meshing out of the cellulose layers. A recent molecular dynamics model proposes that cellulose and pectin layers alternate with each other ^37^. This resembles our observations, with the exception that in the hypothetical model, single-pass cellulose layers alternate with layers of pectic matrix. Our observations do not show alternations of 1-fiber thick layers with meshing, but rather that the cellulose layers can be much thicker (Figure 3E and supplemental video 2) and can vary in thickness. In tomograms proximal to the leading edge of the lamellae, the cell wall adjacent to the plasma membrane displayed increased amounts of meshing (Figure 5D-O) in the form of extensive patches. This is consistent with previous AFM work that identified a layer of “meshwork” interfacing between the plasma membrane and the first layers of cellulose fibers of the cell wall ^26^ that disappeared upon treatment with pectate lyase. Because of the topological nature of the AFM technique, the extent to which this meshwork pervades the cell wall could not be assessed. In our experiments, the meshing is detectable until ∼ 1 μm deep in the cell wall (before the meshing to fiber ratio falls below 0.5, Figure 5H and Figure 6G) as smaller patches and inter-fibrillar tethers (Figure 2 and Figure 4A-C). Such organization is reminiscent of previous Solid State Nuclear Magnetic Resonance (SSNMR) results stating that pectins, when dimethyl-esterified, are cross-linked by calcium, form a branched network that establishes contacts with the cellulose fibers on up to 50% of their surface ^39,40^. Following the amount of meshing throughout a given lamella, we measured a relevant decrease of the quantity of meshing in tomograms from ∼0 to 3.5 μm deep in the cell wall (Figure 5H, supplemental video 3). This gradient of meshing suggests it is secreted and accumulated at the cell wall-PM interface and in the inner-surface layers and gradually drops out of the deeper, older layers of tightly bundled cellulose layers following the same exclusion mechanism explained above (Figure 5C). This confines the meshing at the inner-surface of the cell wall, less than ∼1 μm deep. This model of cell wall build-up implies structurally different Z-regions. In the light of previous data, we can thus link the structure (fiber – meshing connectivity) to the function (stiffness of the cell wall and its ability to stretch) such that different layers of the wall confer different mechanical properties ^39,40^.

The plasma membrane-adjacent, younger layers are comprised of a mix of meshing-interconnecting and less-bundled cellulose fibers conferring more stiffness to these inner-layers, potentially allowing for more reversible elastic deformations and preventing irreversible plastic deformation. The more cell-distal, older layers of the cell wall are comprised mostly of tightly bundled cellulose fibers devoid of meshing. These cellulose bundles are non-pectin-crosslinked, therefore loosening the respective layers of the cell wall and potentially allowing them to stretch and undergo irreversible plastic deformation (creep). Our observations highlight the importance of considering the cell wall as a 3-D polylamellate structure to grasp its mechanical properties fully.

### Treatments affecting the morphology and quantity of meshing suggest the meshing is HG

The visible thinning of the cell wall upon PL treatment (Figure S7D and E) suggests that removal of the major pectin, HG, leads to a collapse of the stacked layers, suggesting that HG acts as a filler, intercalating between the cellulose layers (Figure 2B and E). This is in line with the latest model of how pectins arrange around the cellulose fibers in the onion primary cell wall ^37,40^.

The effect of the PL enzyme on the meshing morphology and abundance (Figure 6G), and the fact that this enzyme specifically digests HGs with low degrees of methyl-esterification indicates that the meshing is predominantly composed of demethylesterified HGs. This is also confirmed by the COS488 homogalacturonan-specific staining patterns upon PL treatments (Figure S6D-F). Moreover, the reticulated networks of purified HGs with a ∼40% methyl-esterified composition have nearly identical morphology (Figure S8) to what was observed *in situ* in native cell walls, adding plausibility to the proposal that HG is the major component of the meshing. This is consistent with previous observations, showing that digestion with the same pectate lyase enzyme leads to the disappearance of the interfacial pectin between the cell wall and the plasma membrane ^26^.

HGs interact strongly with divalent cations like calcium. Specifically calcium ions have been shown to cross-link demethylated HGs, leading to a shift in mechanical properties ^23^. In the light of this, treatment with BAPTA, a divalent cation chelator, was expected to reduce reticulation. In our data, upon BAPTA treatment, we can still see the branched meshing (Figure 6A, C, D, and G), indicating that maintenance of this structure does not depend on calcium concentration and that branching can still occur in low-calcium concentrations.

Overall, our results regarding the nature and distribution of the meshing throughout the cell wall suggest that it consists of demethylesterified HGs. This fits well with the current understanding of pectin biosynthesis where pectins are synthesized in a methylated state in the Golgi Apparatus and demethylated *in muro* by endogenous methylesterases ^24,41,42^ and strongly interact with the cellulose fibers ^39^, participating in the stiffness of the primary cell wall ^40^.

The remnants observed in one instance (Figure 6B, E and F) could be more complex pectic polysaccharides such as rhamno-galacturonan-II, or the hemicellulosic component xyloglucan, shown to coat the cellulose fibers, possibly tethering them together as we observed in our tomograms (Figure 4C) ^27^ or HGs with a higher degree of methylation, and therefore little affected by the enzyme.

### Our measurements of the diameter of the cellulose fibers cannot weight in favor of the 18-or 24-glucan chain model

Averaging of the cellulose fiber diameter in the three conditions considered was performed in an effort to 1) get an idea of the cross-sectional diameter of cellulose fibers *in situ* and 2) check whether PL or BAPTA treatments could have an effect on fiber thickness, as it has been reported that other wall polysaccharides coat the cellulose fibers^27^. Given the difficulty of assessing the width of a density line in cryo-ET because of defocus, we opted for the Full-Width-at-Half-Maximum (FWHM) standardized method (Figure 7D). By this method, we measured fiber diameters ranging from 5.3 to 6.3 nm. PL or BAPTA treatment did not substantially alter the diameter of the cellulose fibers, therefore suggesting that HGs do not coat the cellulose fibers longitudinally. This does not preclude punctual covalent bindings between the HGs and the cellulose fibers, as mentioned in previous models ^43^. Previous reports had measured several hundred fiber diameters in onion walls *in situ* by AFM. The measurements were between 3.5 to 7 nm, which falls within the range of our measurements^26,44^. While an AFM and cryo-EM study favor of the 18 glucan-chain model of an elementary fiber involving CSCs made of hexamers of trimers^11,44^, Solid State NMR indicates the elementary cellulose fiber is composed of 24 glucan chains made by hexamers of tetrameric cellulose synthases^39^. Our measurement, which averages range from ∼5-6 nm, with a maximal resolution of 3.8 nm (Figure S11), do not allow a conclusion one way or the other.

### Limitations of this study

This work represents, to our knowledge, the first report to use cryo-FIB milling followed by cryo-ET to observe the plant cell wall. Despite the high quality of the data achieved in this study, the throughput of the method is limited by the lengthy milling times and the low survival rate of lamellas from milling to tilt series acquisition. Future work will focus on testing other enzymatic treatments, e.g., specifically directed towards hemicelluloses like xyloglucan or combinations of enzymes and assess the impact of the degraded component on the structure of the cell wall in 3-dimensions.

Our study focused on the milling of the periclinal cell wall, as due to their geometry as large flat surfaces (Figure 1E and F), they were more amenable to our approach. This does not preclude the feasibility of milling the anticlinal cell walls, even though initial attempts resulted in unstable, very short lamellae unsuitable for cryo-ET. It would be of high interest to visualize the anticlinal cell walls, notably because they constitute interfaces between neighboring cells of the same scale, allowing the visualization of the pectic mid-lamella ^45^. Subsequent studies focusing on the anticlinal cell wall might be enabled by the introduction of more powerful ion sources which are expected to render milling through thick slabs of material more feasible ^46^. Lastly, while preparing the sample and positioning the cell wall peel on the EM grid, the polarity of the peel relative to the orientation of the onion bulb was not tracked, something to consider for future experiments that would allow comparison of the relative fiber orientation in different scales.

## Methods

### Onion cell wall peel preparation

White onions were purchased the day of or the day prior to the experiments at the local Pavilions supermarket (845 E California Blvd, Pasadena, CA 91106). Peels at the various concentric scales used throughout this work were generated as described in ^28,29^. Briefly, the scales were sliced longitudinally with a sharp knife or razor blade. Then the middle of each slice, where the width is more or less constant, a ∼4 cm long piece was cut out. An incision was made with a sharp razor blade approximately 1 cm away from the edge, creating “handles”. Then these handles were used to pull apart the epidermal layer away from the parenchyma of the scale. This resulted in peels about 1 cm in width and 2-3 cm in length. These peels were incubated for at least 20 min in HEPES buffer (20 mM HEPES, 0.1% Tween-20, pH 6.8 with KOH) and remained in it until freezing. Before freezing, each cell wall peel was mounted between slide and coverslip and screened with a table-top microscope equipped with phase-contrast to ensure that the peel had a homogenous surface of cleanly ruptured cells where only the cell wall remained. Phase-contrast allowed visualization of the remaining floppy, jagged-looking anticlinal cell walls, indicating that peeling of the cell wall was successful.

### Quantification of the aspect ratios of epidermal cells by light microscopy

A large montage of the epidermal cell wall peels was acquired by light microscopy using a Nikon 90i epifluorescence microscope. These maps were segmented with ImageJ with the following method (Figure S9): i) out-of-focus cells and folded-over peels were masked out manually to avoid distorted cell segmentations using the polygon selection tool of imageJ and deleting the selected areas (Figure S9A and B). ii) A binary mask was applied on the montages in order to select the outline of the cells, and the resulting mask was gaussian-filtered (2 pixel) and skeletonized (Figure S9C). iii) The “Analyze particles” tool was used to detect closed cells and calculate their aspect ratio (Figure S9D and E).

### Enzymatic treatments and staining

Pectate lyase from *Aspergillus* (Megazyme, 180 U/mg, Cat # E-PCLYAN2) at 4.7U/mL (8uL of stock solution in 5mL of 50 mM CAPS buffer, pH 10) ^10^ and BAPTA calcium chelation at 2mM (Sigma Aldrich – Cat # A4926) treatments were performed on cell wall peels generated as described above. Treatments were carried out for 3 hours and 10 min, respectively, on the peels.

To screen for the effectivity of the treatments prior to vitrification, staining of non-treated, BAPTA-or PL-treated onion peels by a homogalacturonan-specific probe, Chitosan OligoSaccharide coupled with Alexa-488 (COS488) (Figure S6B) was performed based on a protocol provided by Jozef Mravec (personal communication): 1:1000 dilution from the mother solution kindly provided by Jozef Mravec (kept at −20C wrapped in foil) in 50 mM MES buffer pH 5.8 for 15 min. Peels were then washed with DI water 3 consecutive times before being mounted between a slide and coverslip and then screened by confocal laser scanning microscopy ^47^.

### Purified pectin preparation

Citrus-derived high homogalacturonan content purified pectins were kindly provided by Professor Hans-Ulrich, from Herbstreith & Fox (https://www.herbstreith-fox.de/en/): Pectin Classic CU 701 (38% methyl-esterification, 89% galacturonic acid content). 10mL of 2.5% (w/v) aqueous pectic solutions were made (pH 3.4 according to manufacturer’s MSDS sheet). Serial dilutions at 0.25% and 0.125% were then prepared from the 2.5% solution.

### Plunge-freezing

#### Onion cell wall peels

The cell wall peels previously incubated in HEPES buffer were laid on a slide with a drop of HEPES buffer to keep the cell wall hydrated. After incubation in HEPES buffer, the peels were mounted in a drop of HEPES on a slide. A tangential light was shined at the peel to increase visibility. If possible, a magnifying glass affixed on a support can be used. Small rectangular pieces (∼2 × 3 mm) were cut out of the cell wall peel with a sharp razor blade and carefully dragged on the carbonated side of glow-discharged (15mA – 1min) Quantifoil R2/2 NH2 Cu EM grids (EMSdiasum). Plunge freezing was performed with a 60/40 ratio ethane/propane mix and an FEI Vitrobot Mark IV (Thermo Fischer). Humidity was set at 50%, temperature at 20°C. Grids were first manually backblotted for 6 s in order to attach the cell wall peel firmly to the carbon, followed by two autoblottings (front and back) 5 s, maximal blot force (25) and a drain time of 3 s.

#### Purified pectins

5uL of 2.5%, 0.25% and 0.125% purified pectin was pipetted onto Quantifoil R2/2 NH2 Cu EM grids (EMSdiasum) and the grids were plunge frozen at 100% humidity, 20C with a blot time of 4 s, a medium blot force of 10 and a drain time of 1 s.

### Cryo-FIB milling

During the grid clipping stage, prior to milling, orientation of the peel is important, so the long side of the rectangle was positioned parallel to the notch in autogrid holders (Thermo Fisher) machined with a notch. Like this, the shorter side of the anticlinal cell walls are orthogonal to the FIB beam leading to less obstructed areas of the periclinal cell wall and thus more potential FIB-milling targets. Autogrids were placed in a shuttle and inserted into a Versa 3D dual-beam FIB/SEM microscope with a field emission gun (FEG) (FEI) equipped with a PP3000T cryo-transfer apparatus (Quorum Technologies). They were maintained at −175°C at all times by a cryo-stage ^48^. To reduce sample charging and protect the sample from curtaining during milling, the grids were sputter-coated with platinum at 15mA for 60 s. Thin lamellae were generated with the gallium ion beam at 30 kV at angles ranging from 10 to 17°. Rough milling was done at high currents, ranging from 0.3 nA to 100 pA, until the lamellae measured 1 μm in thickness under the FIB view. The current was then progressively brought down to 10 pA for the final milling steps until the measured thickness was between 100 and 200 nm Final polishing by tilting the sample 0.5 to 1° to homogenize the lamella thickness was also done at 10 pA. During the whole procedure, imaging with the SEM beam was done at 5 kV and 27 pA. SEM overviews were used to precisely outline and measure the respective aspect ratios (width vs. length) of the milled cells. When In-chamber Gas Injection System (GIS) Pt coating was performed, the needle was set at 26C and flushed for ∼10s before injection onto the onion peel. The injection was performed for ∼5 s at a distance of +2 mm from eucentric height.

### Confocal microscopy

Confocal analysis of the onion cell wall peels stained with the COS488 stain was performed on a ZEISS LSM880 equipped with Airy Scan and a GaAsP detector. Magnification used was 40x (C-Apochromat 40x/1.2 W Korr M27). Channel settings were set as follows and kept constant throughout the conditions screened: For the Alexa 488 channel the excitation Ar laser (488 nm) was set to 0.3% power, the gain was set to ∼700 and pinhole was set to ∼10 AU with a pixel dwell time of ∼2 μs. A GaAsP detector was used, and the detection range was set from 499 to 630 and the 488 main beam splitter was used. Trans-channel was set with a gain of ∼450. Z-stacks were acquired with the optimal Z-step defined by the software, 1.55 μm.

### Electron-cryotomography

Tilt-series acquisition was performed on a Titan Krios (Thermo Fisher) equipped with a GIF post-column energy filter (Gatan) and a K3 direct detector 6k × 4k (Gatan). Data acquisition was controlled via SerialEM ^49^ with a 3° tilt increment for a total range of ±60° or ±50°, a defocus of −10 μm, and a total dose up to 80 e^-^/A^2^. No pre-tilt was applied and a bi-directional tilt scheme was used. Tilt series were then aligned via patch tracking with the IMOD package ^50^ reconstructed using weighted back projection and the SIRT-like filter set to 15 iterations.

### Mapping out tomograms on the milled cells

Orientation of the grid is lost during the transfer from the cryo-SEM chamber to the cryo-TEM autoloader. The grid can be rotated and/or flipped over. This necessitated correlating the orientations found in cryo-SEM and the cryo-TEM data. We, therefore, used the high-resolution montage maps of the lamellae as the reference where the different fields of view of the tomograms can be seen (Figure S10A, blue rectangles). Using Adobe Illustrator, the high-resolution TEM montages were correlated with the TEM grid montages and the cryo-SEM-overviews of the lamellae. The latter are flipped and rotated if needed to fit the final orientation in the TEM used for data collection (Figure S10B). Finally, the angle between the X-axis of the tomograms and the long axis of the cell was registered (Figure S10C, blue and red lines, respectively). This ensured the precise knowledge of the long axis of the milled cell within each tomogram (Figure S10D, black dashed line), which in turn allowed the extraction of biologically relevant numbers. Depth of the tomograms in the cell wall was computed using the nominal milling angle as the inclination and the projected distance d between the leading edge of the lamella (top of the lamella, identified by the presence of platinum) and the center of the ROIs for tilt series acquisition (Figure 1J).

### Sub-tomogram averaging and cross-sectional measurements

Sub-tomogram extraction, alignment, and averaging were performed using the Dynamo software package^51^. Initial orientations and positions of cellulose fibers segments were determined using geometrical tools for particle picking in Dynamo^52^. Regions of the filaments with minimal bending and overlapping were traced in 4x binned tomograms. Centers of the particles were placed every ∼70 Å along the filament. Final sub-volumes were extracted from 2x binned tomograms with a final pixel size of 6.7 Å and 40×40×40 box size. The total number of sub-tomograms ranged from 750 to 1100 for all three datasets. Initial reference for particle alignment was generated by averaging segments with azimuth randomized orientations. Iterative alignment and averaging procedures were performed according to gold-standard in Dynamo. A loose cylindrical mask was applied for the alignment step. The final mask corrected FSC was estimated in RELION3 using a soft-edge mask (Figure S11)^53^.

### Measurement of the cross-sectional thickness

The *sideview-profile-average* script^54^ was used by tracing an open contour in the middle of the fiber in 3dmod. The following parameters were used: step 1 pixel, length 30 pixels, and thickness 10 pixels. The output json files were imported into R. The average pixel intensities were double normalized relative to the lowest and highest pixel values in each profile to compare curves between conditions. The Full-Width-at-Half-Maximum (FWHM) was used by measuring the width of the gaussian bell at 0.5 relative pixel intensity.

### Tomogram segmentation

#### Fiber segmentation

Segmentation was performed on filtered tomograms with the default parameters of EMAN2 (low-pass gaussian cutoff of 0.25 and high-pass gaussian cutoff of 5px) and Convolutional Neural Networks (CNN) ^30^ were used to recognize the fibers in the tomograms (Figure S1A-C). Training was performed on several tomograms by boxing ∼20 positive examples and ∼100 negative examples. The positive examples were precisely segmented using a graphical tablet (*Wacom Cintiq 21uX*) and the CNNs were trained with the default parameters except for the learn rate that was increased in some instances to 0.001 instead of the default 0.0001. The outcome of the trained CNN was checked on the boxed particles and if satisfactory the CNN was applied on the tomogram. Eventually, a second round of training was performed with additional boxes from another tomogram from the same dataset or on itself. The resulting CNN map was then carefully examined versus the filtered tomogram to ensure they agreed, and segmentation was specific to the fibers. For tomograms acquired over the same session on the same lamellae, the same CNN was able to generalize well and segment accurately. Tomograms from different datasets and different lamellae usually required retraining a CNN.

Satisfactory CNN segmented volumes were then transferred into *Amira* (Thermo Fisher) to perform template matching fiber tracing with the *TraceX Amira* plugin ^31^ (Figure S1D-F) in order to model the fibers as a set of connected nodes. To be able to optimize parameters, we reduced the processing time by binning twice (binning 8 total) the CNN maps. The first step, *Cylinder Correlation*, was performed with the following starting parameters: cylinder length of 50 pixels, an angular sampling of 5, and missing wedge compensation was toggled. The diameter of the template (outer cylinder radius) was set to closely match the apparent thickness of the fibers in the tomogram, usually 4 pixels. As advised in the Amira user guide section 3.8 on the *XTracing Extension*, the mask cylinder radius was set to 125% of the outer cylinder radius. The outcome was visually checked to see if the fibers were detected correctly and not too many artefacts were generated. Parameters were slightly modified one-by-one if needed to improve the output. The subsequent step, *Trace Correlation Lines* was performed with the following nominal parameters: minimal line length 60 pixels, direction coefficient 0.3, and minimal distance of 2-times outer-cylinder diameter used previously. Minimum seed correlation and minimum correlation are tomogram-dependent parameters. These values were defined on the correlation field by defining the reasonable correlation value range. The minimum seed correlation and minimum continuation quality are the upper and lower limits of the range, respectively. For the search cone, length was set to 80, angle to 37°, and minimal step size was 10%. The outcome was visually checked to see if the fibers were being traced correctly. To do so, we used the *Spatial Graph View* function and checked for artificial fiber trackings. Parameters were modified if needed to enhance fiber detection and reduce false discoveries. Because of the inherent nature of the signal of cryo-ET volumes and their CNN maps, punctate signals would generate and propagate artefactual vertical (parallel to the Z-dimension) lines. These were first selected by using a Tensor XZ and Tensor ZZ visualizer in the *Spatial Graph View* window and identifying the appropriate thresholds. After the coordinates of all fibers were extracted as a .xml file, fiber tracks with values above/below the thresholds were trimmed out.

#### Meshing segmentation and quantification

The method to output the CNN maps recognizing the meshing is identical to the one used to segment the fibers. We were unable to generate a CNN that could specifically pick up the meshing. Instead, we resorted to training CNNs that could recognize all features in the tomograms and then subtracted this density map with the one generated from the fiber-trained CNN. This allowed isolation of identified features that were not fibers, assuming that everything that is not fiber belongs to the “meshing” feature. This was done using a custom script called *MeshingSubtract* (available upon demand) that relies on IMOD and bash commands. First, the fiber-CNN map was thresholded. The level of the threshold is chosen in order to mask the fibers as accurately as possible. This mask is then subtracted from the meshing-CNN map to create the subtracted meshing map.

To quantify the volume occupancy of these two features, the *imodauto* command was used. A threshold of 0 was used on the masked fiber-CNN. For the subtracted meshing map, the threshold was chosen in order to segment as accurately as possible the meshing by comparing with the low-pass filtered tomogram. Both resulting segmentations were joined using the *imodjoin* command and the *imodinfo* command was used to compute the volume occupancy of each segmented feature (the value taken was the cylinder volume).

### Data extraction

Point (containing only point number, x-, y- and z-coordinates) and segment data (containing only point numbers) from the Amira-Avizo (Thermo Fisher) software was exported as tab-delimited files. The reformat_amira_output.m ^55^, available from https://schurlab.ist.ac.at/downloads/ was used to convert these files into IMOD formatted tab-delimited text files, which were then further analyzed using custom scripts in python.

First, all contours with an out-of-plane angle of larger than 70 degrees were removed, as those did not correspond to fibers but rather to tomogram reconstruction artifacts. For each model, the long axis of the cell was accurately determined as detailed above. Then the model was rotated around the y-axis to reposition the volume according to the angle applied during the milling step and lost during the volume flattening occurring during tomogram reconstruction.

To overcome the uneven spacing of points on contours exported from Amira, fiber contours were interpolated using cubic splines resulting in a sampling rate of 1 nm along the length of the fiber. From these reoriented volumes in the cell wall, fiber length, radius of curvature, slope of the fiber, clockwise angle of the fiber relative to the cell’s long axis,

The length of individual fibers was calculated as the sum of distances between neighboring points along its run:

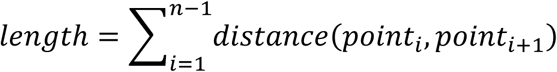

With n being the number of points of the given contour representing the fiber.

The curvature radius was calculated by averaging the local curvature radii over all triples of neighboring points within a given contour. For this the reciprocal relationship between the Menger curvature and the curvature radius was employed:

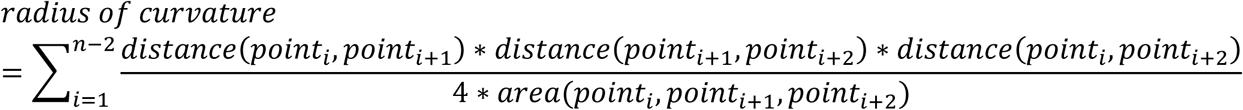

With n being the number of points of the given contour representing the fiber.

Prior to calculating local slopes along the run of a fiber, the sequence of the points was, if necessary, adjusted so that the first point of the contour would have a lower y-coordinate than the last point of the contour. This was done to establish a common direction for all fibers within a tomogram. To calculate the slope between two neighboring points on a contour representing a fiber the difference between their z-coordinate is divided by their distance in the xy-plane:

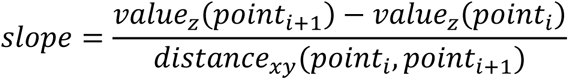

With value_z_(point) extracting the z-coordinate of a given point and distance_xz_(point, point) calculating the distance between two points only considering x- and y-coordinates.

For calculating the average z-height of a fiber, the z-coordinates of all points in the respective contour were averaged.

For calculating the angle between a fiber and the long axis of the cell, the orientation of the fiber was approximated by a vector pointing from its end with the lower y-coordinate value to its end with the higher y-coordinate value. The vector representing the long axis of the cell was calculated from the orientation of the cell on the grid and the rotations applied during tomogram reconstruction.

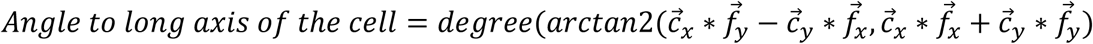

With x and y representing the x and y scalars of the vectors of the long axis of the cell 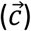 or the fiber 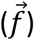, respectively. The resulting angles in radians were then transformed to degrees as depicted in the figures.

### Data analysis and visualization

All the data analysis, data exploration and statistical analysis was performed with R. Specifically, statistical analysis of the value distributions and modelling was done by the mixed models method using the *dipTest* and *mixtools* R packages.

## Acknowledgments

This work was supported by the Howard Hughes Medical Institute (HHMI) and grant R35 GM122588 to Grant J Jensen, and Austrian Science Fund (FWF): P33367 to Florian KM Schur. We thank Noé Cochetel for his GST4 guidance and great help in data analysis, discovery and representation with the R software. We thank Hans-Ulrich Endress for graciously providing us with the purified citrus pectin, Jozed Mravec for generating and providing the COS488 probe. Cryo-EM work was done in the Beckman Institute Resource Center for Transmission Electron Microscopy at Caltech.

This article is subject to HHMI’s Open Access to Publications policy. HHMI lab heads have previously granted a nonexclusive CC BY 4.0 license to the public and a sublicensable license to HHMI in their research articles. Pursuant to those licenses, the author-accepted manuscript of this article can be made freely available under a CC BY 4.0 license immediately upon publication.

## Bibliography

1. Bar-On, Y. M., Phillips, R. & Milo, R. The biomass distribution on Earth. Proc. Natl. Acad. Sci. 115, 6506–6511 (2018).

2. Johnson, M. P. Photosynthesis. Essays Biochem. 60, 255 (2016).

3. Verbančič, J., Lunn, J. E., Stitt, M. & Persson, S. Carbon Supply and the Regulation of Cell Wall Synthesis. Molecular Plant vol. 11 75–94 (2018).

4. Voragen, A. G. J., Coenen, G. J., Verhoef, R. P. & Schols, H. A. Pectin, a versatile polysaccharide present in plant cell walls. Struct. Chem. 20, 263–275 (2009).

5. Zhang, B., Gao, Y., Zhang, L. & Zhou, Y. The plant cell wall: Biosynthesis, construction, and functions. Journal of Integrative Plant Biology vol. 63 251–272 (2021).

6. Ruel, K., Nishiyama, Y. & Joseleau, J.-P. Crystalline and amorphous cellulose in the secondary walls of Arabidopsis. Plant Sci. 193–194, 48–61 (2012).

7. Makarem, M. et al. Distinguishing Mesoscale Polar Order (Unidirectional vs Bidirectional) of Cellulose Microfibrils in Plant Cell Walls Using Sum Frequency Generation Spectroscopy. J. Phys. Chem. B 124, 8071–8081 (2020).

8. Cosgrove, D. J. Nanoscale structure, mechanics and growth of epidermal cell walls. Current Opinion in Plant Biology vol. 46 77–86 (2018).

9. Wang, X. & Cosgrove, D. J. Pectin methylesterase selectively softens the onion epidermal wall 1 yet reduces acid-induced creep 2 3 4 Running title: Effects of pectin methylesterase on cell wall mechanics. J. Exp. Bot. (2020).

10. Zhang, T., Tang, H., Vavylonis, D. & Cosgrove, D. J. Disentangling loosening from softening: insights into primary cell wall structure. Plant J. 100, 1101–1117 (2019).

11. Purushotham, P., Ho, R. & Zimmer, J. Architecture of a catalytically active homotrimeric plant cellulose synthase complex. Science (2020).

12. Nixon, B. T. et al. Comparative Structural and Computational Analysis Supports Eighteen Cellulose Synthases in the Plant Cellulose Synthesis Complex. Sci. Rep. 6, 28696 (2016).

13. Li, S., Lei, L., Somerville, C. R. & Gu, Y. Cellulose synthase interactive protein 1 (CSI1) mediates the intimate relationship between cellulose microfibrils and cortical microtubules. Plant Signal. Behav. 7, 1–5 (2012).

14. Chan, J., Coen, E., Chan, J. & Coen, E. Interaction between Autonomous and Microtubule Guidance Systems Controls Cellulose Synthase Report Interaction between Autonomous and Microtubule Guidance Systems Controls Cellulose Synthase Trajectories. Curr. Biol. 1–7 (2020).

15. Diotallevi, F. & Mulder, B. The cellulose synthase complex: A polymerization driven supramolecular motor. Biophys. J. 92, 2666–2673 (2007).

16. Kuki, H. et al. Quantitative confocal imaging method for analyzing cellulose dynamics during cell wall regeneration in Arabidopsis mesophyll protoplasts. Plant Direct 1, e00021 (2017).

17. Wang, Y. & Jiao, Y. Cellulose Microfibril-Mediated Directional Plant Cell Expansion: Gas and Brake. Mol. Plant 13, 1670–1672 (2020).

18. Anderson, C. T., Carroll, A., Akhmetova, L. & Somerville, C. Real-time imaging of cellulose reorientation during cell wall expansion in Arabidopsis roots. Plant Physiol. 152, 787–796 (2010).

19. Aouar, L., Chebli, Y. & Geitmann, A. Morphogenesis of complex plant cell shapes: The mechanical role of crystalline cellulose in growing pollen tubes. Sex. Plant Reprod. 23, 15–27 (2010).

20. Sampathkumar, A. et al. Subcellular and supracellular mechanical stress prescribes cytoskeleton behavior in Arabidopsis cotyledon pavement cells. Elife 3, e01967 (2014).

21. Park, Y. B. & Cosgrove, D. J. A revised architecture of primary cell walls based on biomechanical changes induced by substrate-specific endoglucanases. Plant Physiol. 158, 1933–1943 (2012).

22. Zhang, T., Vavylonis, D., Durachko, D. M. & Cosgrove, D. J. Nanoscale movements of cellulose microfibrils in primary cell walls. Nat. Plants 3, 17056 (2017).

23. Cao, L., Lu, W., Mata, A., Nishinari, K. & Fang, Y. Egg-box model-based gelation of alginate and pectin: A review. Carbohydrate Polymers vol. 242 116389 (2020).

24. Shin, Y., Chane, A., Jung, M. & Lee, Y. Recent Advances in Understanding the Roles of Pectin as an Active Participant in Plant Signaling Networks. Plants 10, 1712 (2021).

25. Bidhendi, A. J. & Geitmann, A. Geometrical Details Matter for Mechanical Modeling of Cell Morphogenesis. Dev. Cell 50, 117-125.e2 (2019).

26. Zhang, T., Zheng, Y. & Cosgrove, D. J. Spatial organization of cellulose microfibrils and matrix polysaccharides in primary plant cell walls as imaged by multichannel atomic force microscopy. Plant J. 85, 179–192 (2016).

27. Zheng, Y., Wang, X., Chen, Y., Wagner, E. & Cosgrove, D. J. Xyloglucan in the primary cell wall: assessment by FESEM, selective enzyme digestions and nanogold affinity tags. Plant J. 93, 211– 226 (2018).

28. Kafle, K. et al. Cellulose microfibril orientation in onion (Allium cepa L.) epidermis studied by atomic force microscopy (AFM) and vibrational sum frequency generation (SFG) spectroscopy. Cellulose 21, 1075–1086 (2013).

29. Durachko, D., Park, Y. B., Zhang, T. & Cosgrove, D. Biomechanical Characterization of Onion Epidermal Cell Walls. BIO-PROTOCOL 7, (2017).

30. Chen, M. et al. Convolutional neural networks for automated annotation of cellular cryo-electron tomograms. Nat. Methods 14, 983–985 (2017).

31. Rigort, A. et al. Automated segmentation of electron tomograms for a quantitative description of actin filament networks. J. Struct. Biol. 177, 135–144 (2012).

32. Wilson, L. A., Deligey, F., Wang, T. & Cosgrove, D. J. Saccharide analysis of onion outer epidermal walls. Biotechnol. Biofuels 14, (2021).

33. Sampathkumar, A. et al. Patterning and Lifetime of Plasma Membrane-Localized Cellulose Synthase Is Dependent on Actin Organization in Arabidopsis Interphase Cells. Plant Physiol. 162, 675–688 (2013).

34. Sánchez-Rodríguez, C. et al. BRASSINOSTEROID INSENSITIVE2 negatively regulates cellulose synthesis in Arabidopsis by phosphorylating cellulose synthase 1. Proc. Natl. Acad. Sci. 114, 3533– 3538 (2017).

35. Speicher, T. L., Li, P. Z. & Wallace, I. S. Phosphoregulation of the Plant Cellulose Synthase Complex and Cellulose Synthase-Like Proteins. Plants 7, (2018).

36. Djafari Petroudy, S. R. Physical and mechanical properties of natural fibers. in Advanced High Strength Natural Fibre Composites in Construction 59–83 (Woodhead Publishing, 2017).

37. Zhang, Y., Yu, J., Wang, X., Durachko, D. M. & Zhang, S. Molecular insights into the complex mechanics of plant epidermal cell walls. Science (80-.). 372, 35 (2021).

38. Nicolas, W. J., Ghosal, D., Tocheva, E. I., Meyerowitz, E. M. & Jensen, G. J. Structure of the bacterial cellulose ribbon and its assembly-guiding cytoskeleton by electron cryotomography. J. Bacteriol. (2020).

39. Wang, T. & Hong, M. Solid-state NMR investigations of cellulose structure and interactions with matrix polysaccharides in plant primary cell walls. J. Exp. Bot. 67, 503–514 (2016).

40. Phyo, P., Gu, Y. & Hong, M. Impact of acidic pH on plant cell wall polysaccharide structure and dynamics: insights into the mechanism of acid growth in plants from solid-state NMR. Cellulose 26, 291–304 (2019).

41. Braybrook, S. A. & Peaucelle, A. Mechano-Chemical Aspects of Organ Formation in Arabidopsis thaliana: The Relationship between Auxin and Pectin. PLoS One 8, e57813 (2013).

42. Goubet, F. & Mohnen, D. Subcellular localization and topology of homogalacturonan methyltransferase in suspension-cultured Nicotiana tabacum cells. Planta 1999 2091 209, 112– 117 (1999).

43. Cosgrove, D. J. Re-constructing our models of cellulose and primary cell wall assembly. Current Opinion in Plant Biology vol. 22 122–131 (2014).

44. Song, B., Zhao, S., Shen, W., Collings, C. & Ding, S.-Y. Direct Measurement of Plant Cellulose Microfibril and Bundles in Native Cell Walls. Front. Plant Sci. 11, 479 (2020).

45. Zamil, M. S. & Geitmann, A. The middle lamella—more than a glue. Phys. Biol. 14, 015004 (2017).

46. McClelland, J. J. et al. Bright focused ion beam sources based on laser-cooled atoms. Appl. Phys. Rev. 3, 011302 (2016).

47. Mravec, J. et al. Tracking developmentally regulated post-synthetic processing of homogalacturonan and chitin using reciprocal oligosaccharide probes. Dev. 141, 4841–4850 (2014).

48. Rigort, A. et al. Micromachining tools and correlative approaches for cellular cryo-electron tomography. J. Struct. Biol. 172, 169–179 (2010).

49. Mastronarde, D. N. Automated electron microscope tomography using robust prediction of specimen movements. J. Struct. Biol. 152, 36–51 (2005).

50. Kremer, J. R., Mastronarde, D. N. & McIntosh, J. R. Computer visualization of three-dimensional image data using IMOD. J. Struct. Biol. 116, 71–6 (1996).

51. Castaño-Díez, D., Kudryashev, M., Arheit, M. & Stahlberg, H. Dynamo: A flexible, user-friendly development tool for subtomogram averaging of cryo-EM data in high-performance computing environments. J. Struct. Biol. 178, 139–151 (2012).

52. Castaño-Díez, D., Kudryashev, M. & Stahlberg, H. Dynamo Catalogue: Geometrical tools and data management for particle picking in subtomogram averaging of cryo-electron tomograms. J. Struct. Biol. 197, 135–144 (2017).

53. Zivanov, J. et al. New tools for automated high-resolution cryo-EM structure determination in RELION-3. Elife 7, (2018).

54. Ortega, D. R. et al. Repurposing a chemosensory macromolecular machine. Nat. Commun. 11, 1– 13 (2020).

55. Dimchev, G., Amiri, B., Fäßler, F., Falcke, M. & Schur, F. K. Computational toolbox for ultrastructural quantitative analysis of filament networks in cryo-ET data. bioRxiv 2021.05.25.445599 (2021).

