## Supplemental figures for "Bimodally oriented cellulose fibers and reticulated homogalacturonan networks - A direct visualization of Allium cepa primary cell walls"

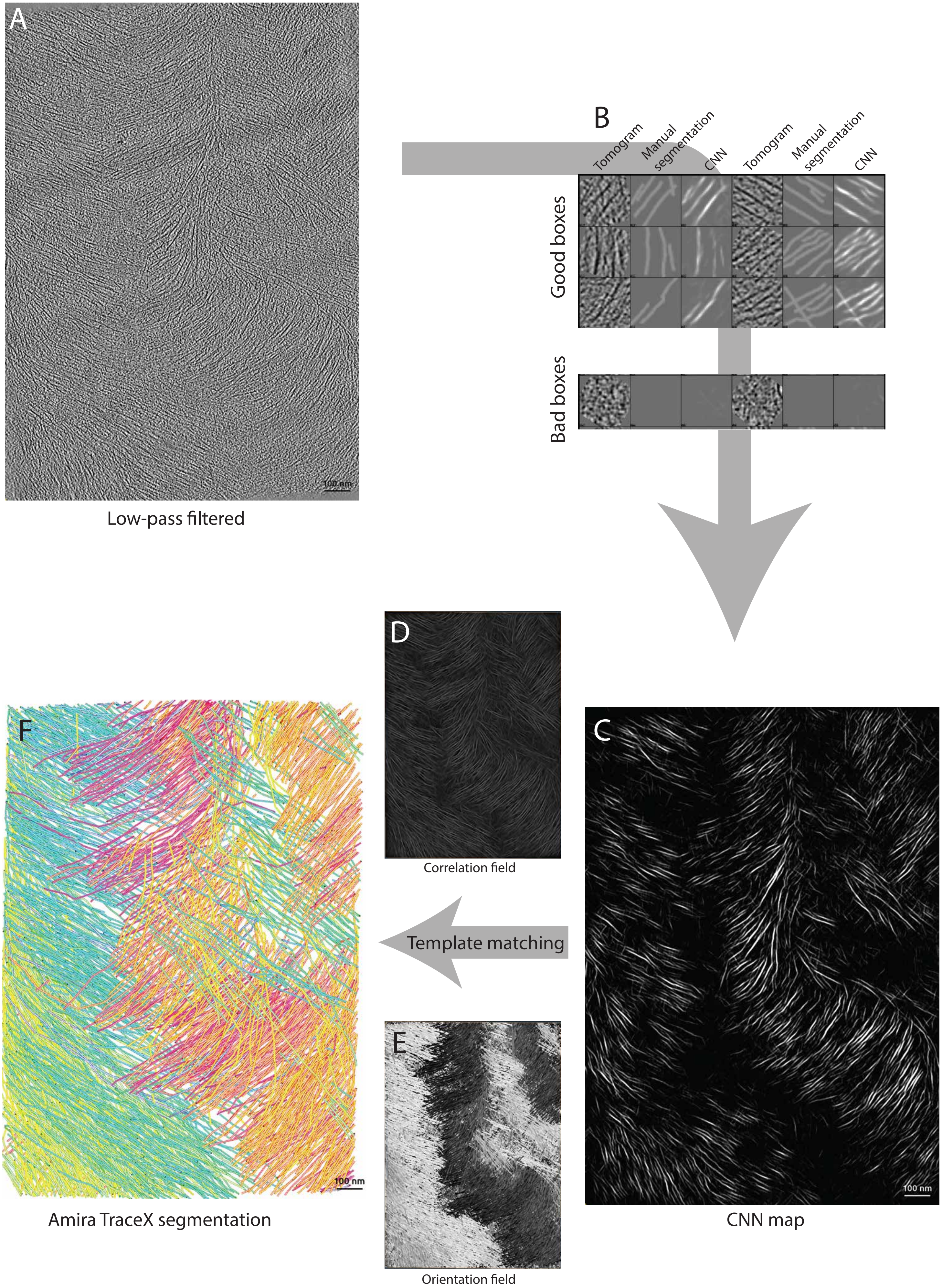
A
100 nm
Low-pass filtered
B
Tomogram
Manual segmentation
CNN
Tomogram
Manual segmentation
CNN
Good boxes
Bad boxes
C
100 nm
CNN map
D
Correlation field
Template matching
E
Orientation field
F
100 nm
Amira TraceX segmentation

**Supplemental figure 1 | Convolutional Neural Networks and Template matching based segmentation of the tomograms**

(**A**) Example tomographic slice of a low-pass filtered tomogram. (**B**) Example training boxes of cellulose fibers. From left to right, the columns are the boxed sub-tomograms, the manual segmentation provided for the training, and the CNN segmentation. (**C**) EMAN2-CNN segmented tomogram. (**D, E**) Correlation and orientation field outputs generated by Amira during the template matching step with the template being a 50 pixels long, 4 pixels wide outer-cylinder diameter. (**F**) Final segmentation of the segmented fibers.

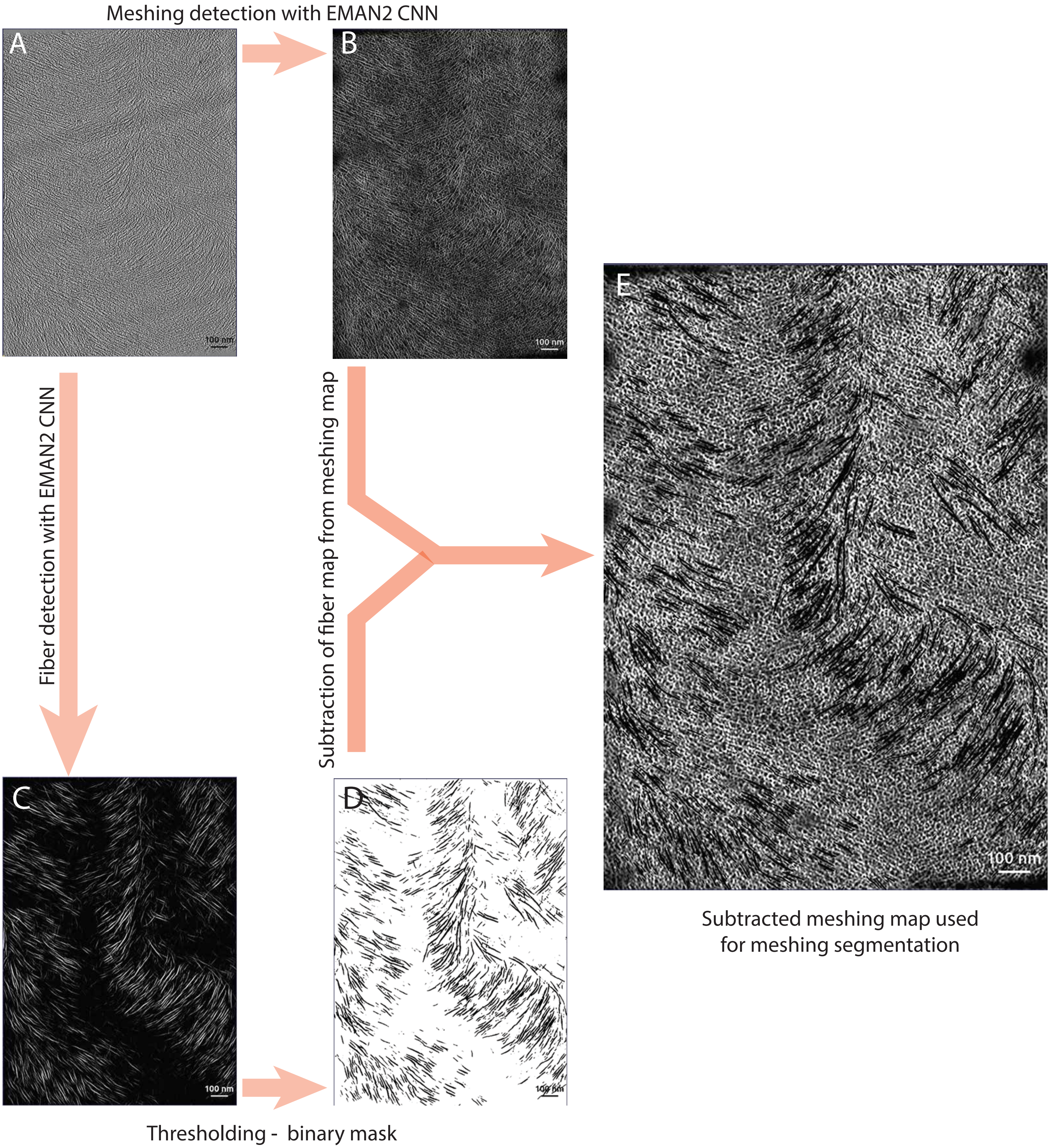
Meshing detection with EMAN2 CNN
A
B
100 nm
100 nm
Fiber detection with EMAN2 CNN
Subtraction of fiber map from meshing map
E
100 nm
C
D
100 nm
Thresholding - binary mask
Subtracted meshing map used
for meshing segmentation

**Supplemental figure 2 | Segmentation of the meshing**

(**A**) Tomographic slice of a low-pass filtered tomogram. (**B**) Tomogram segmented by a CNN trained to recognize the meshing. (**C**) Tomogram segmented by a CNN trained to recognize the fibers. (**D**) Masked fiber-segmented tomogram. (**E**) Meshing-segmented tomogram (B) subtracted by the thresholded fiber-segmented tomogram (D).

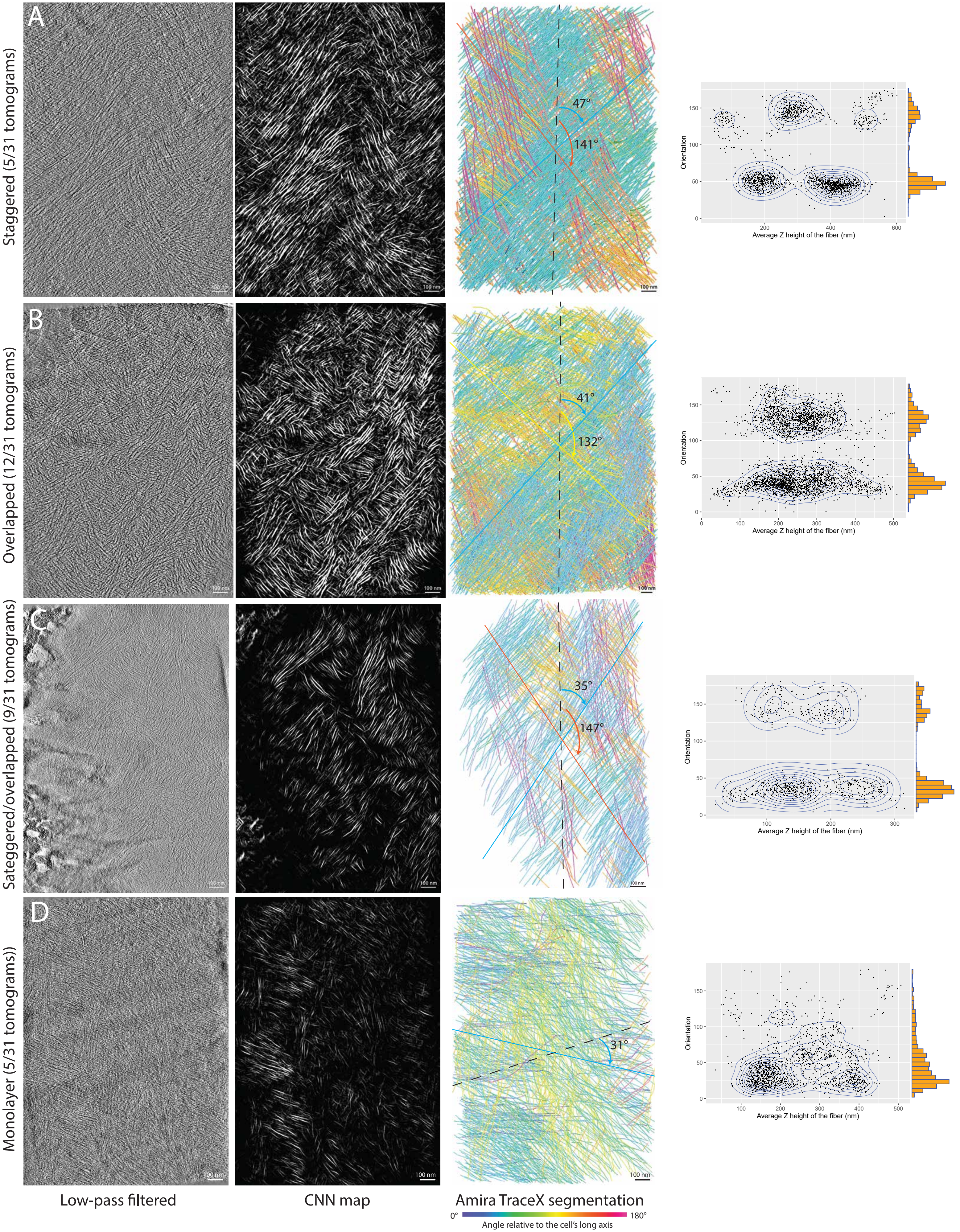

A
B
C
D
Staggered (5/31 tomograms)
Overlapped (12/31 tomograms)
Sateggered/overlapped (9/31 tomograms)
Monolayer (5/31 tomograms))
47°
141°
41°
132°
35°
147°
31°
100 nm
Orientation
Average Z height of the fiber (nm)
Low-pass filtered
CNN map
Amira TraceX segmentation
0°
180°
Angle relative to the cell's long axis

**Supplemental figure 3| The different fiber orientation layering patterns**

From left to right: The low-pass filtered tomograms, the CNN segmented tomograms, the Amira segmented volumes displaying the cell's long axis (black dashed line) and the main modes, and the scatterplot of the orientation of the fiber as a function of its average Z-height. (**A**) The staggered pattern characterized by clean successive -/+ 45° cellulose fiber layers, with clearly defined clusters in the scatterplots. (**B**) The overlapped pattern where the two -/+ 45° are intercalated with each other. (**C**) The overlapped-staggered pattern, similar to (A) but the scatterplot shows an overlapping cluster. (**D**) The monolayer pattern showing only one main mode but a very scattered cluster, as shown in the scatterplots.

### Effect of depth in the cell wall on the angular distribution of fibers

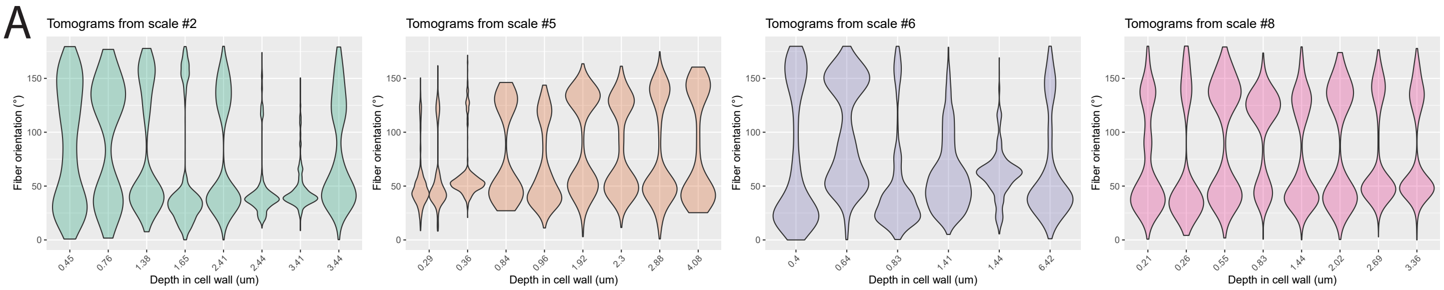

### Effect of aspect ratio of the cell on the angular distribution of fibers

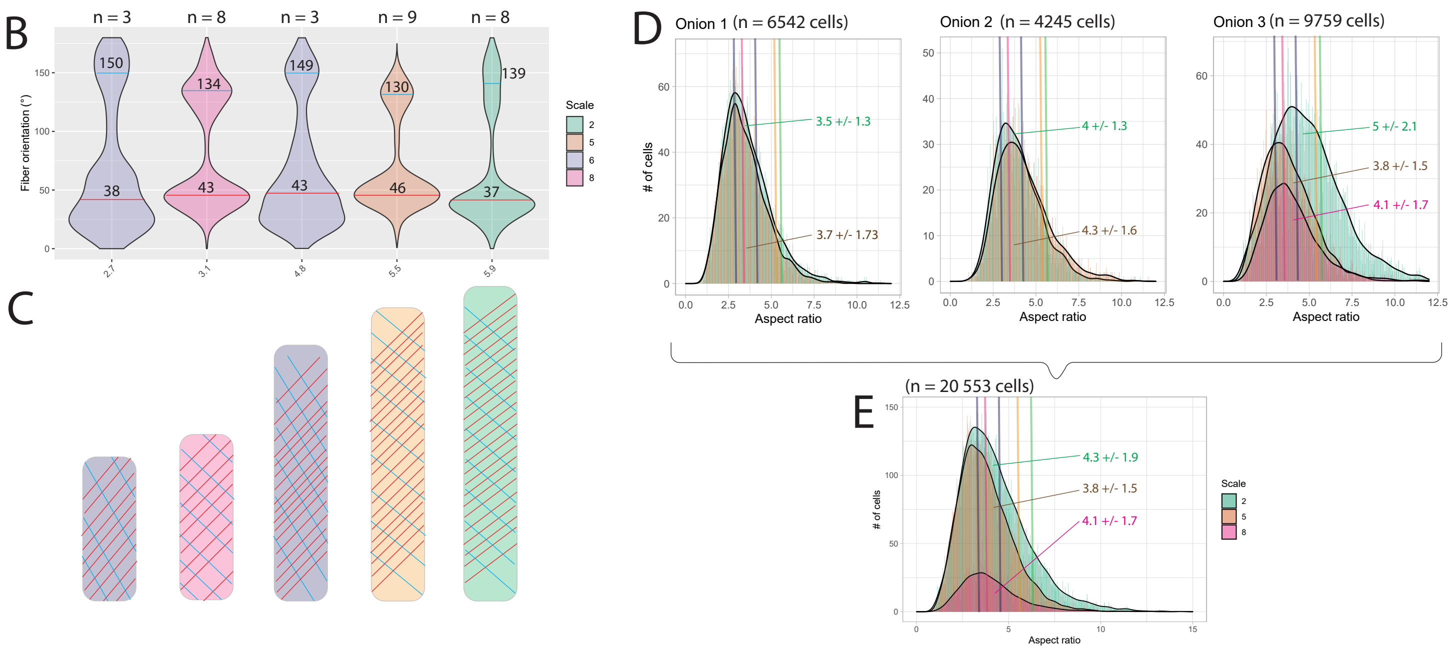

**Supplemental figure 4 | The bimodal angular pattern is found in cells of all aspect ratios and at all depths** (**A**) Violin plots for each scale (scale #2, 5, 6 and 8, from left to right), showing the distribution of the fiber angles as a function of the depth of the tomographic volume in the milled cell wall. One violin represents one tomogram. (**B**) Violin plots for each milled cell, showing the distribution of the fiber angles as a function of the aspect ratio of the milled cell. The violins are color-coded per scale and the X-axis shows the aspect ratios of the milled cells. (**C**) Drawings of cells with the corresponding aspect ratios to the cells milled shown in (B). (**D**) Distribution of the aspect ratios of cells screened by light microscopy (see methods) in 3 different onions. Colored vertical lines represent the aspect ratios of the milled cells in (B) and show that the milled cells fall within the range of aspect ratios of their respective scales. (**E**) Same as (D) but all cells from the three onions were pooled.

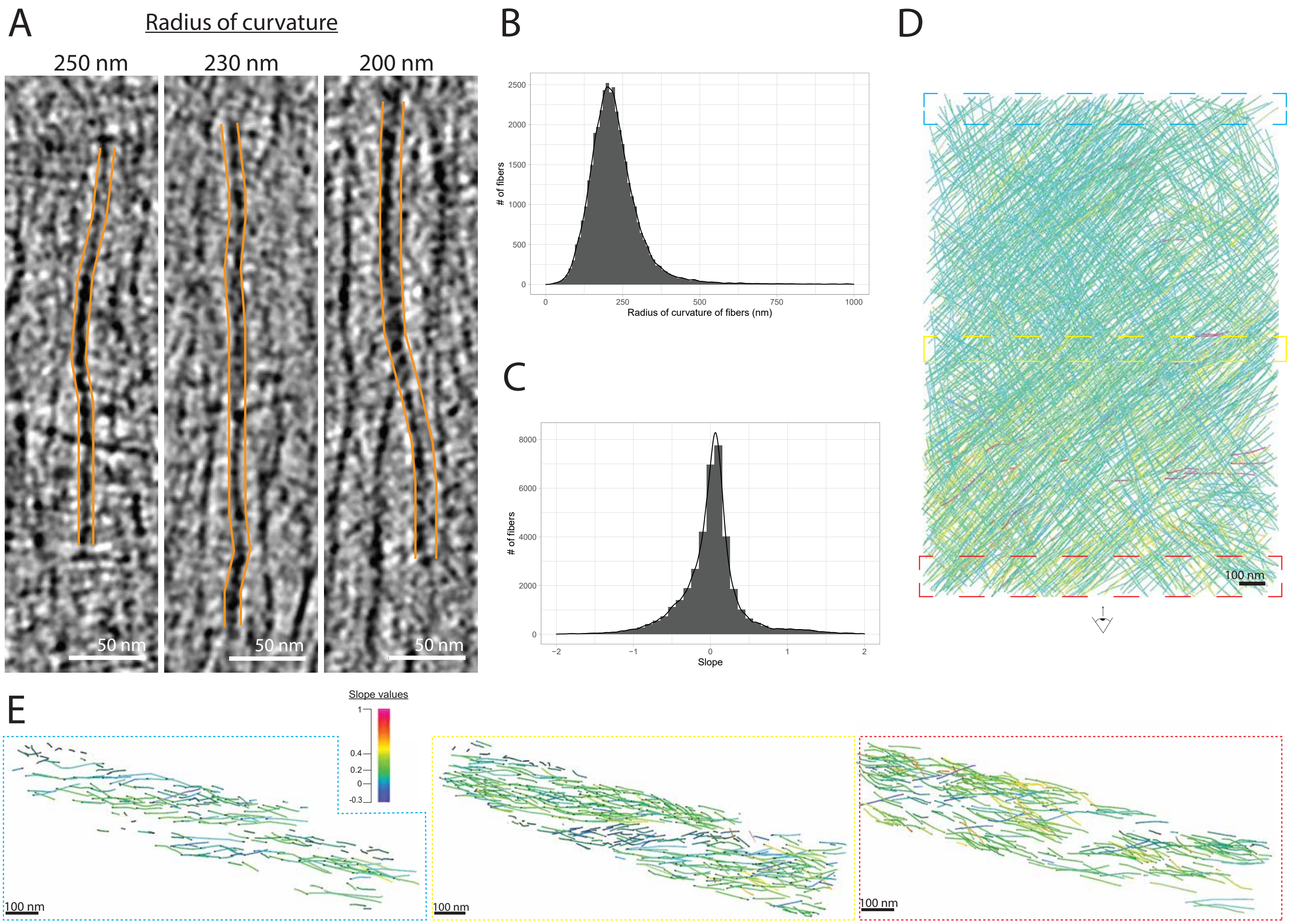

**Supplemental figure 5 | The fibers travel straight and horizontally in the cell wall**

(**A**) Examples of fibers with average radii of curvature spanning a range of 250 to 200nm. The fibers are highlighted in orange. (**B**) Global distribution of the average radius of curvature of all fibers across all scales, with an average at 225 ± 90 nm. (**C**) Global distribution of the average slopes of the fibers across all scales, with an average at 0.02 ± 0.4. (**D**) Segmented tomographic volume with color coding reflecting the slope of the fibers. Cyan fibers hold a slope value around 0. (**E**) Transversal views of small sub-volumes (boxed with the corresponding color in (D)) showing how the fibers are relative to the horizontal after lamella angle correction.

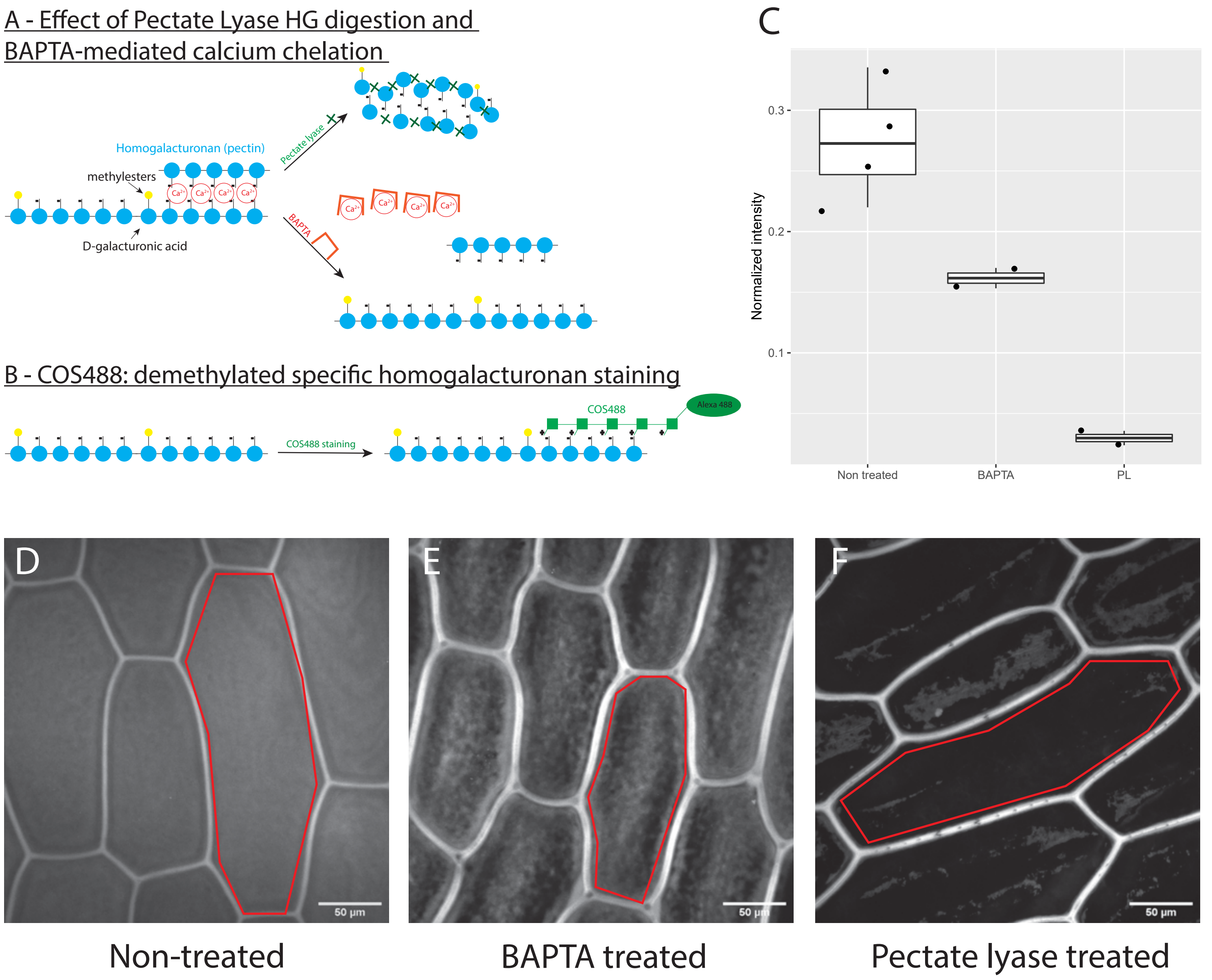

**Supplemental figure 6 | COS staining and effect of PL and BAPTA on HG pectins**

(**A**) Homogalacturonans demethylated groups can be cross-linked by the calcium present in the cell wall. Pectate lyase will sever the connections between the D-galacturonic acid units (top). BAPTA chelates the calcium, preventing cross-linking of pectins (bottom). (**B**) COS-488 is used to stain demethylated HGs by specifically binding to the demethylated groups. (**C**) Fluorescence intensity quantification of the effect of the PL and BAPTA treatments on the COS-488 stained peels. (**D**) Non-treated COS-488 stained peels showing homogeneous staining of the periclinal cell wall. (**E**) BAPTA-treated COS-488 stained peels showing a different, heterogeneous staining in the periclinal cell wall. (**F**) PL-treated COS-488 stained peels showing a significant decrease in the intensity of the signal in the periclinal cell wall.

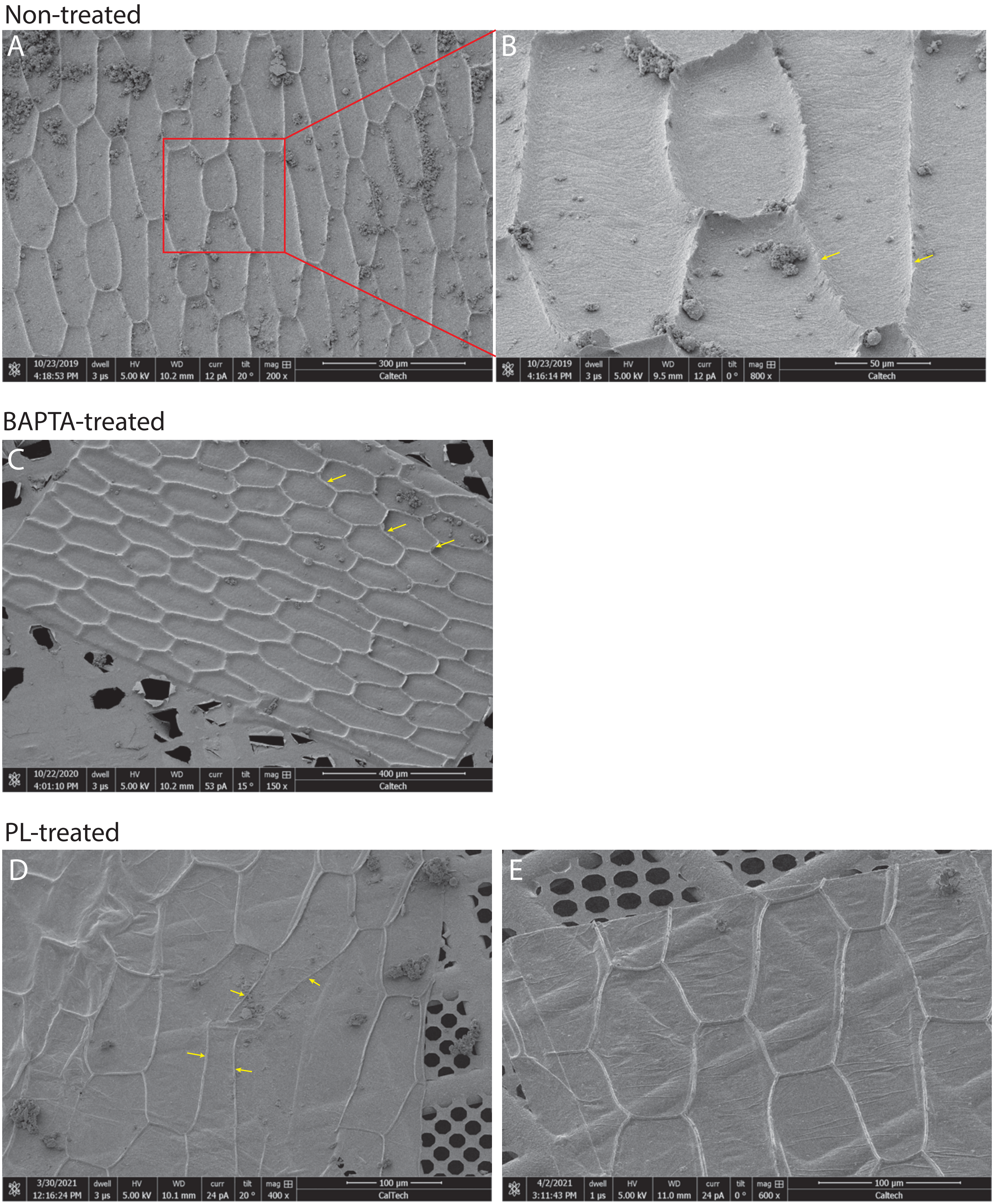
Non-treated
A
B
10/23/2019 4:18:53 PM dwell 3 μs HV 5.00 kV WD 10.2 mm curr 12 pA tilt 20 ° mag 200 x 300 μm Caltech
10/23/2019 4:16:14 PM dwell 3 μs HV 5.00 kV WD 9.5 mm curr 12 pA tilt 0 ° mag 800 x 50 μm Caltech
BAPTA-treated
C
10/22/2020 4:01:10 PM dwell 3 μs HV 5.00 kV WD 10.2 mm curr 53 pA tilt 15 ° mag 150 x 400 μm Caltech
PL-treated
D
E
3/30/2021 12:16:24 PM dwell 3 μs HV 5.00 kV WD 10.1 mm curr 24 pA tilt 20 ° mag 400 x 100 μm CalTech
4/2/2021 3:11:43 PM dwell 1 μs HV 5.00 kV WD 11.0 mm curr 24 pA tilt 0 ° mag 600 x 100 μm Caltech

**Supplemental figure 7 | Effect of PL and BAPTA treatment on the morphology of the onion cell wall peels**

(**A**) SEM overview of a non-treated cell wall peel. (**B**) Magnified view of red rectangle in (A) showing the details of the periclinal cell wall and the anticlinal cell wall remains (yellow arrows). (**C**) SEM overview of a BAPTA-treated cell wall peel with a similar morphology to the non-treated cell wall peel. Yellow arrows point out to the anticlinal cell wall remains. (**D**) SEM overview of a PL-treated cell wall peel showing a thinned-out periclinal cell wall and detached anticlinal remains (yellow arrows). (**E**) Additional example of a PL-treated cell wall peel showing how thin they are compared to non-treated and BAPTA-treated peels, as the topology of the grid bars and the Quantifoil carbon is visible underneath the peel.

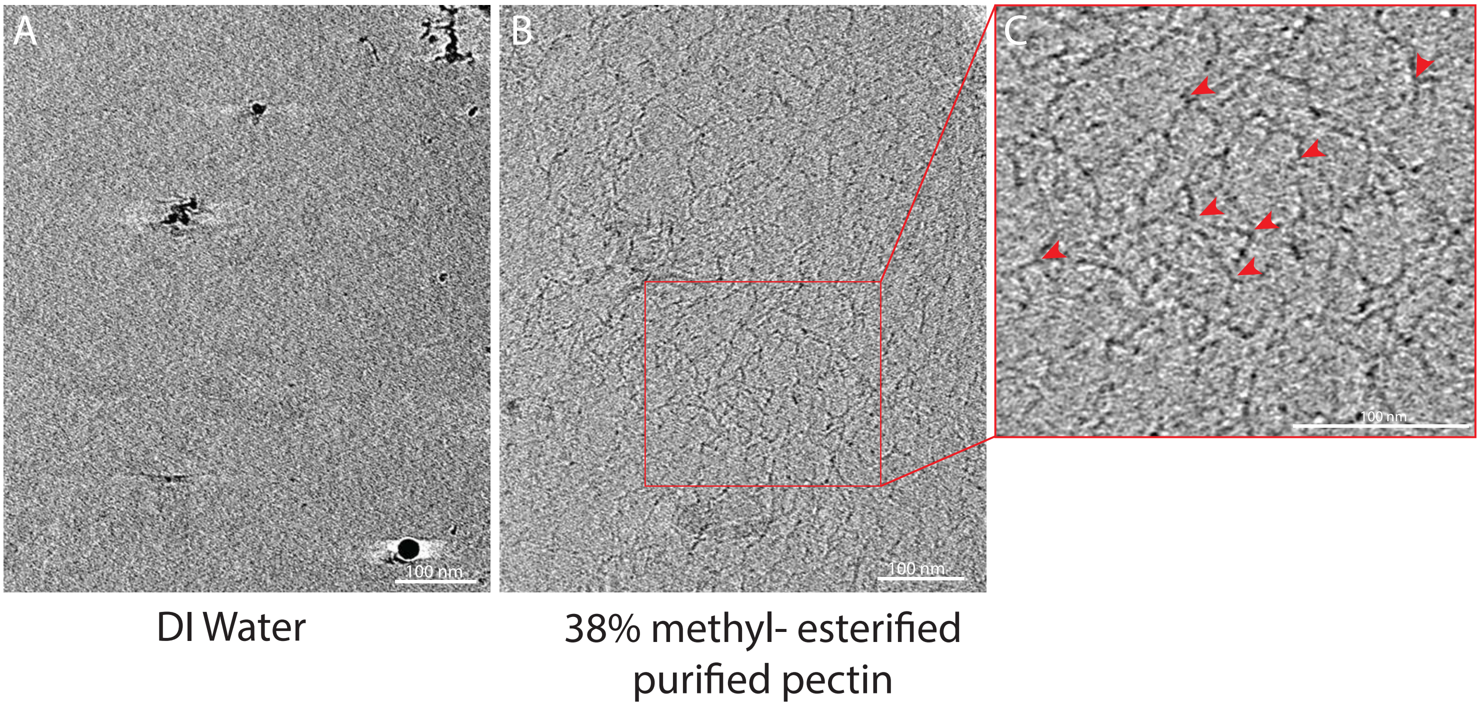

**Supplemental figure 8 | Purified pectins form reticulated meshes on grids**
(**A**) Central tomographic slice from a tomogram of the distilled water used to dissolve pectins (negative control). (**B**) Central tomographic slice from a tomogram of a 0.25% purified citrus pectin with 38% of methyl-esterification, showing a reticulated network. (**C**) Magnified view from (B) showing details of the pectin meshing in the form of short, branched segments (red arrowheads).

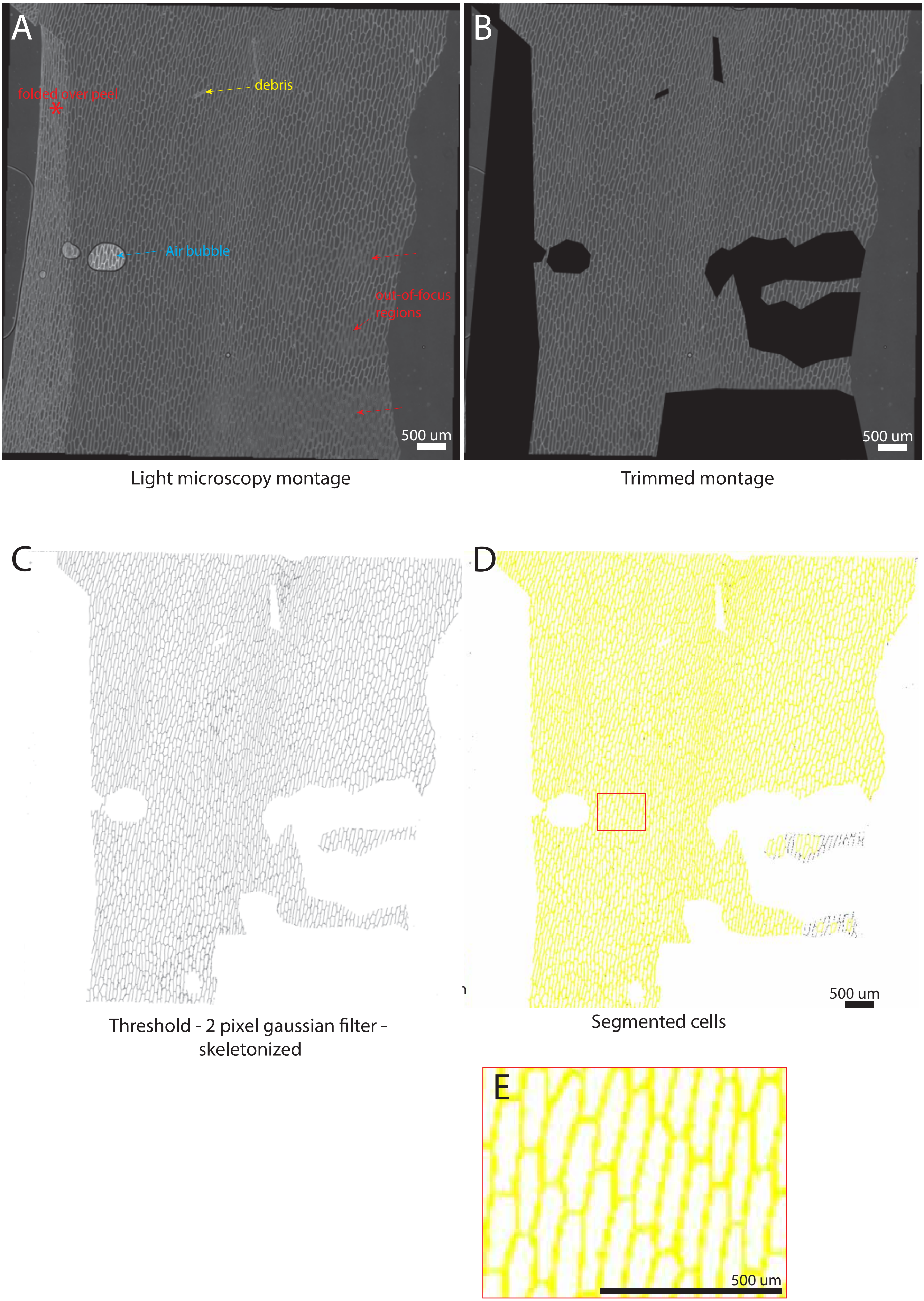

A
folded over peel
debris
Air bubble
out-of-focus regions
500 um
B
500 um
Light microscopy montage
Trimmed montage
C
D
500 um
Threshold - 2 pixel gaussian filter - skeletonized
Segmented cells
E
500 um

**Supplemental figure 9 | Segmentation of the cells in light microscope montages of cell wall peels**

(**A**) Raw cell wall peel montage showing artefacts such as out-of-focus regions (red arrows), folded parts of the peel (red asterisk), debris (yellow arrow), and air bubbles (blue arrow). (**B**) masked out version of the montage in (A), where all the artefacts were manually removed. (**C**) Thresholded masked montage (B) and gaussian filtered + skeletonized map. (**D**) Cells were segmented using the “Analyze particles” function of ImageJ. (**E**) Magnified view from red rectangle in (D) showing the segmented cells used for the quantification of the aspect ratio of cells.

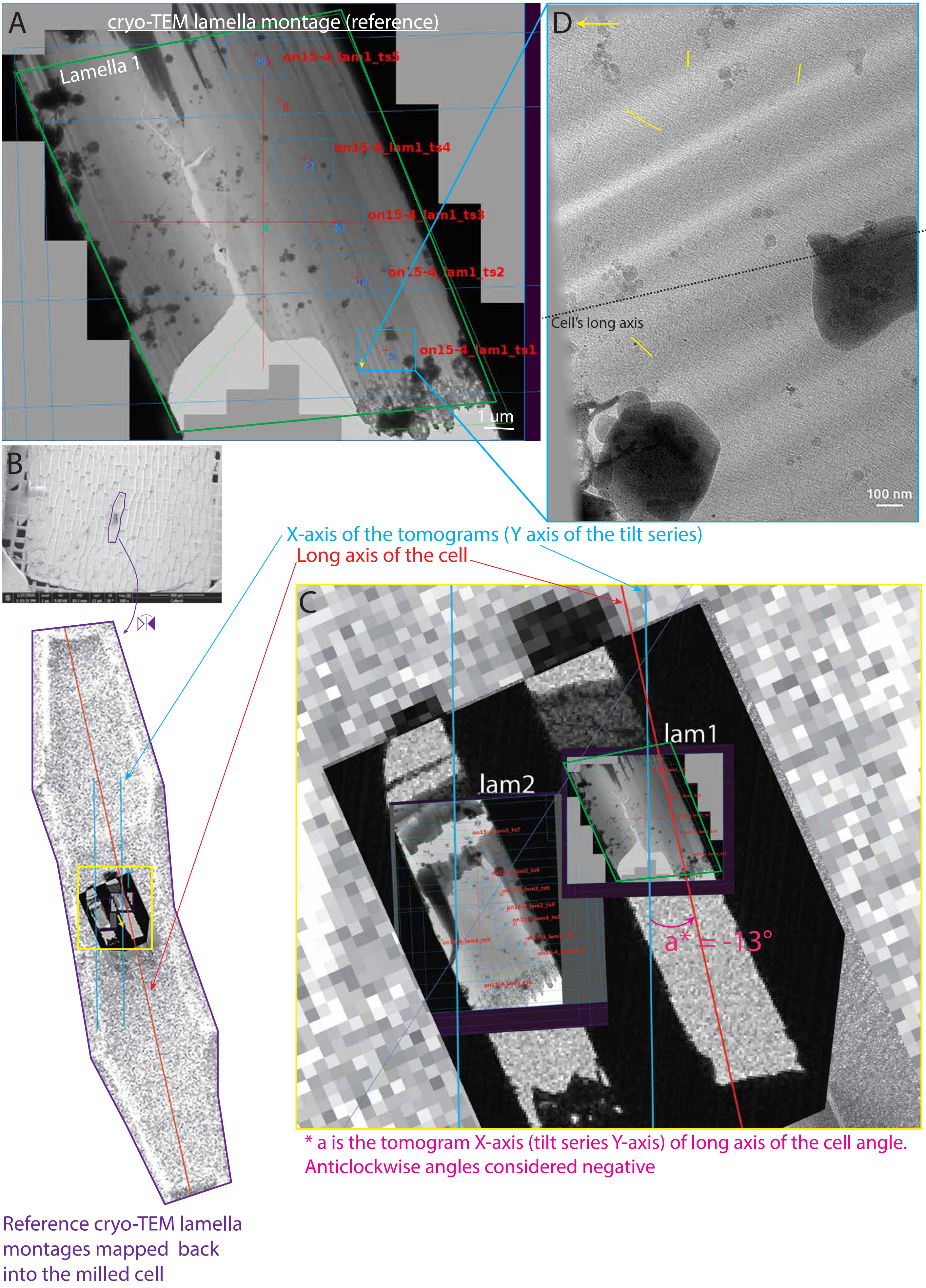
A
cryo-TEM lamella montage (reference)
Lamella 1
on15-4_lam1_ts5
on15-4_lam1_ts4
on15-4_lam1_ts3
on15-4_lam1_ts2
on15-4_lam1_ts1
1 um
B
D
Cell's long axis
100 nm
X-axis of the tomograms (Y axis of the tilt series)
Long axis of the cell
C
lam1
lam2
a* = -13°
* a is the tomogram X-axis (tilt series Y-axis) of long axis of the cell angle.
Anticlockwise angles considered negative
Reference cryo-TEM lamella montages mapped back
into the milled cell

**Supplemental figure 10 | Repositioning of the tomograms within the milled cells**

(**A**) TEM high-resolution montage of the lamella used as reference. Blue rectangles are the ROIs where tilt series were acquired. Short side of the rectangles are the Y-axis of the tilt series and X-axis of the tomogram (90° clockwise rotation). (**B**) Top, SEM overview of an onion peel with the two milled lamellae. Purple outline delimitates the milled cell. Below, magnified and mirrored view of the milled cell outlined above with the reference TEM lamella montage mapped on it. Red and blue lines are the long axis of the milled cell and the X-axis of the tilt series, respectively. (**C**) Magnified view (yellow rectangle in (B)) of the correlated cryo-SEM overview and TEM reference montage of the two lamellae mapped back on the SEM overview. Red and blue lines are the long axis of the milled cell and the X-axis of the tilt series, respectively. The angle between these two lines (a) represents the clockwise angle between the X-axis of the tomograms and the cell's long axis. (**D**) 0° projection showing numerous fibers (a few are outline in yellow). Black dashed line represents the cell's long axis. Yellow arrow represents where the leading edge of the lamella is. The Image is 90° clockwise rotated relative to the blue rectangle in (A).

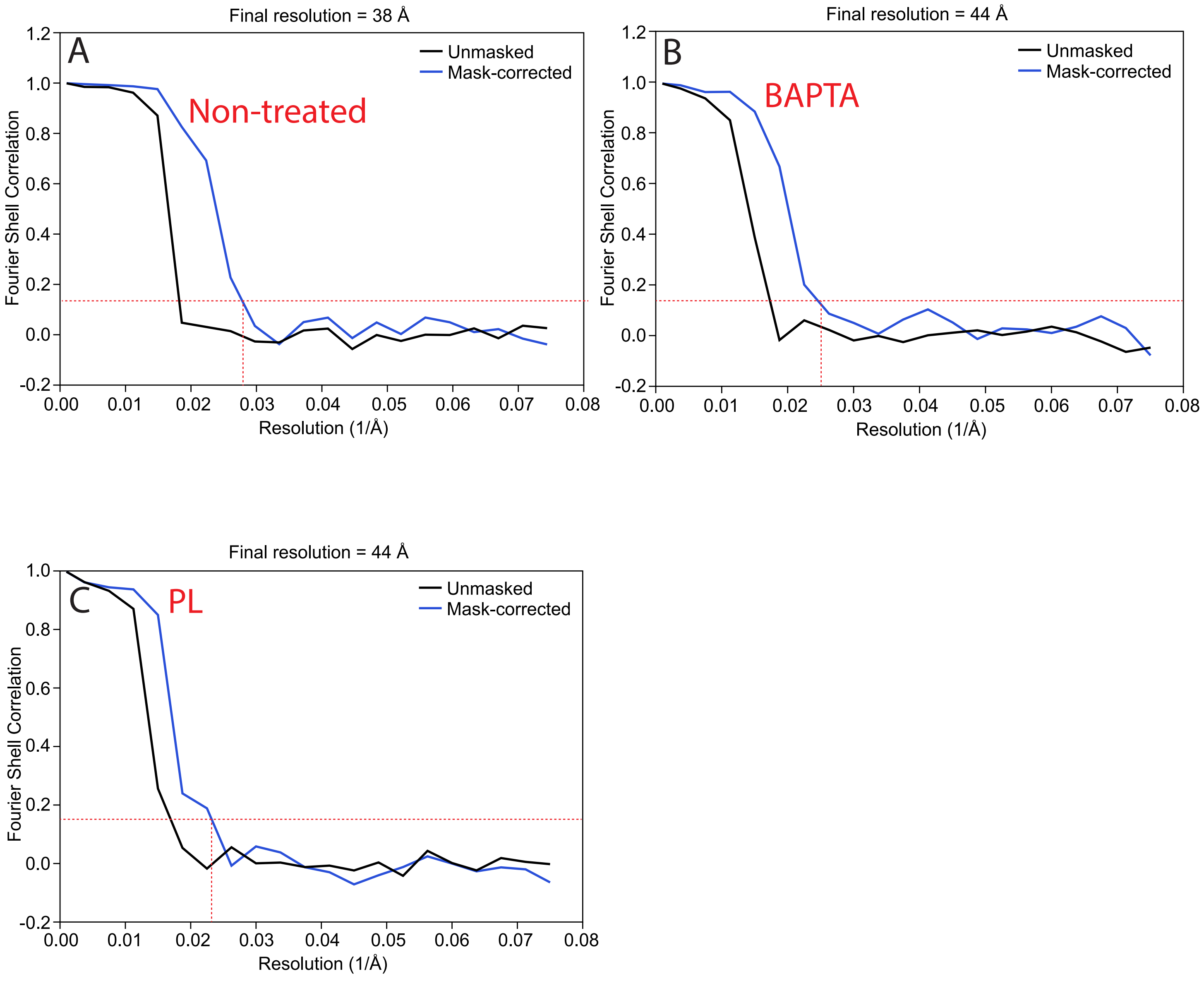

**Supplemental figure 11 | Fourier Shell Correlations of the fiber averages**

(**A-C**) Fourier Shell Correlation curves (unmasked in black and mask-corrected in blue). Red horizontal line is the 0.143 FSC gold standard value and the red vertical line indicates the frequency corresponding to this FSC value when mask-corrected.

| Condition | Individual onion # (freezing time) | Scale | Tomograms |
|---|---|---|---|
| Non-treated | - | - | 31 |
| | 6 (January 2021) | 2 | 8 |
| | 2 (January 2020) | 5 | 9 |
| | 1 (December 2018) | 6 | 6 |
| | 3 (February 2020) | 8 | 8 |
| Pectate lyase-treated | - | - | 6 |
| | 5 (November 2020) | 5 | 5 |
| | 5 (November 2020) | 6 | 1 |
| BAPTA-treated | - | - | 7 |
| | 4 (August 2020) | 7 | 7 |
| Total | 6 onions over the course of 2 years | 5 | 44 |

**Supplemental table 1 | Summary of the tilt-series collected**

Break down of the tilt-series collected by condition (1$^{st}$ column), provenance (2$^{nd}$ column, onions were numbered from 1 to 6) and scale where the lamellae were milled (3$^{rd}$ columns).

#### Supplemental video legends

**Supplemental video 1 | Cryo-FIB milling and cryo-ET of onion cell wall peels**

Cryo-FIB milling allows to generate lamellae in the epidermal onion cell wall peels. Cryo-ET on these lamellae allowed the visualization in 3-dimensions of the cellulose fibers (yellow) and the meshing (red).

**Supplemental video 2 |The bimodal angular pattern of the cellulose fiber orientations in the onion cell wall peels**

The cellulose fibers in the tomograms cluster by their orientation relative to the cell's long axis, creating horizontal layers with following a bimodal angular pattern.

**Supplemental video 3 | The meshing is found in increased amounts proximal to the leading edge of the lamella, i.e. at the surface of the cell wall.**

Tomograms taken close to the top of the cell wall are enriched in meshing in comparison to tomograms taken deeper in the cell wall where the cellulose fibers appear to be more ordered with less meshing present.
